# Sorghum pan-genome explores the functional utility to accelerate the genetic gain

**DOI:** 10.1101/2021.02.02.429137

**Authors:** Pradeep Ruperao, Nepolean Thirunavukkarasu, Prasad Gandham, Sivasubramani S., Govindaraj M, Baloua Nebie, Eric Manyasa, Rajeev Gupta, Roma Rani Das, Harish Gandhi, David Edwards, Santosh P. Deshpande, Abhishek Rathore

## Abstract

Sorghum (*Sorghum bicolor* L.) is one of the most important food crops in the arid and rainfed production ecologies. It is a part of resilient farming and is projected as a smart crop to overcome the food and nutritional challenges in the developing world. The development and characterisation of the sorghum pan-genome will provide insight into genome diversity and functionality, supporting sorghum improvement. We built a sorghum pan-genome using reference genomes as well as 354 genetically diverse sorghum accessions belonging to different races. We explored the structural and functional characteristics of the pan-genome and explain its utility in supporting genetic gain. The newly-developed pan-genome has a total of 35,719 genes, a core genome of 16,821 genes and an average of 32,795 genes in each cultivar. The variable genes are enriched with environment responsive genes and classify the sorghum accessions according to their race. We show that 53% of genes display presence-absence variation, and some of these variable genes are predicted to be functionally associated with drought traits. Using more than two million SNPs from the pan-genome, association analysis identified 398 SNPs significantly associated with important agronomic traits, of which, 92 were in genes. Drought gene expression analysis identified 1,788 genes that are functionally linked to different conditions, of which 79 were absent from the reference genome assembly. This study provides comprehensive genomic diversity resources in sorghum which can be used in genome assisted crop improvement.

## Introduction

Sorghum (*Sorghum bicolor*) is a multi-utility cereal of global importance, and a major food crop in sub-Saharan Africa and South Asia (Ritter *et al*., 2007), (Motlhaodi *et al*., 2014). It is typically a diploid species (*2n=20*) with an estimated genome size of the ∼800Mb sequence (Price *et al*., 2005). It provides important primary and secondary products, such as food, fodder, starch, fiber, biofuels, alcohol, dextrose syrup as well as other products. It is domesticated and further bred for diverse agro-climatic conditions (Li *et al*., 2010) and shows wide diversity at the genome level (Hart *et al*., 2001; Kong *et al*., 2000).

A draft sorghum genome assembly of 730Mb was initially prepared for *Sorghum bicolor* Moench (Paterson *et al*., 2009), followed by an improved assembly of 732.2Mb, covering approximately 91.5% of the genome (McCormick *et al*., 2018). Recently, a sorghum reference genome assembly for the ‘Rio’ line was generated comprising 729Mb (Cooper *et al*., 2019). Each of these genome assemblies is limited to its respective accession and does not reflect the diversity of genes in this species.

The presence or absence of genes or genomic regions between individuals is an important form of genomic variation in plants, and genes can be categorized into core and variable within the species (Golicz, Batley, *et al*., 2016; Saxena *et al*., 2014). The collection of these core and variable genes is known as pan-genome. A study of the pan-genome from a large number of individuals enhances the understanding of species diversity, domestication and breeding history, and provides a more complete characterization of species genes content diversity as demonstrated in rice (Wang *et al*., 2018) and tomato (Gao *et al*., 2019).

Several approaches are available to construct a pan-genome (Golicz, Batley, *et al*., 2016). The classical approach of whole-genome assembly of all genotypes was initially implemented in bacteria, and later developments led to the complementary method to “iteratively map and assemble”, the unmapped sequence reads, demonstrated in *B. oleracea* (Golicz, Bayer, Barker, Edger, H. Kim, *et al*., 2016), *B. napus* (Hurgobin *et al*., 2018), bread wheat (Montenegro *et al*., 2017) and pigeon pea (Zhao *et al*., 2020). The whole genome assembly and comparison method has the advantage in that it can place almost all individual specific genes in a genomic context, but suffers from the inability to distinguish assembly or annotation errors from true biological variation (Bayer *et al*., 2017). It is also unsuitable for large population studies due to the expense of sequencing, assembling and comparing large numbers of genomes. In contrast, the iterative assembly approach can cost effectively assess large numbers of individuals for gene presence/absence variation and hence identify genes that may be relatively rare in a population and not samples in whole genome assembly approaches, though without additional long read data, it is unable to place many of the newly identified genes. Hence the iterative assembly method is most suited for large population diversity studies.

Hence, we assembled a pan-genome using reference and re-sequenced genomes for genetically diverse race-specific sorghum accessions. The sorghum pan-genome was initiated with the reference genome obtained from JGI on Phytozome (McCormick *et al*., 2018), followed by adding to this reference with novel genome sequences from 176 sorghum lines.

We provided structural and potential functional aspects of this pan-genome in the form of genes, single nucleotide polymorphism (SNP) and gene presence and absence variations (PAV). The utility of the pan-genome was demonstrated by identifying candidate functional genes using publicly available SNP chip data, genome-wide association studies and gene-expression assays. These sorghum pan-genome resources will be useful for achieving the sustainable development goals in developing countries by accelerating the genetic gain in arid and semi-arid ecologies.

## Results

### Pan-genome assembly

Genome sequence data with minimum 10X coverage from earlier studies (Guo *et al*., 2019; Valluru *et al*., 2019) were used for pan-genome assembly (Supplementary Table 1). The pan-genome was constructed using 176 sorghum accessions using an iterative mapping and assembly approach, similar to Brassica (Golicz, Bayer, Barker, Edger, H. Kim, *et al*., 2016) and pigeon pea (Zhao *et al*., 2020). (Figure 1).

**Figure 1:**
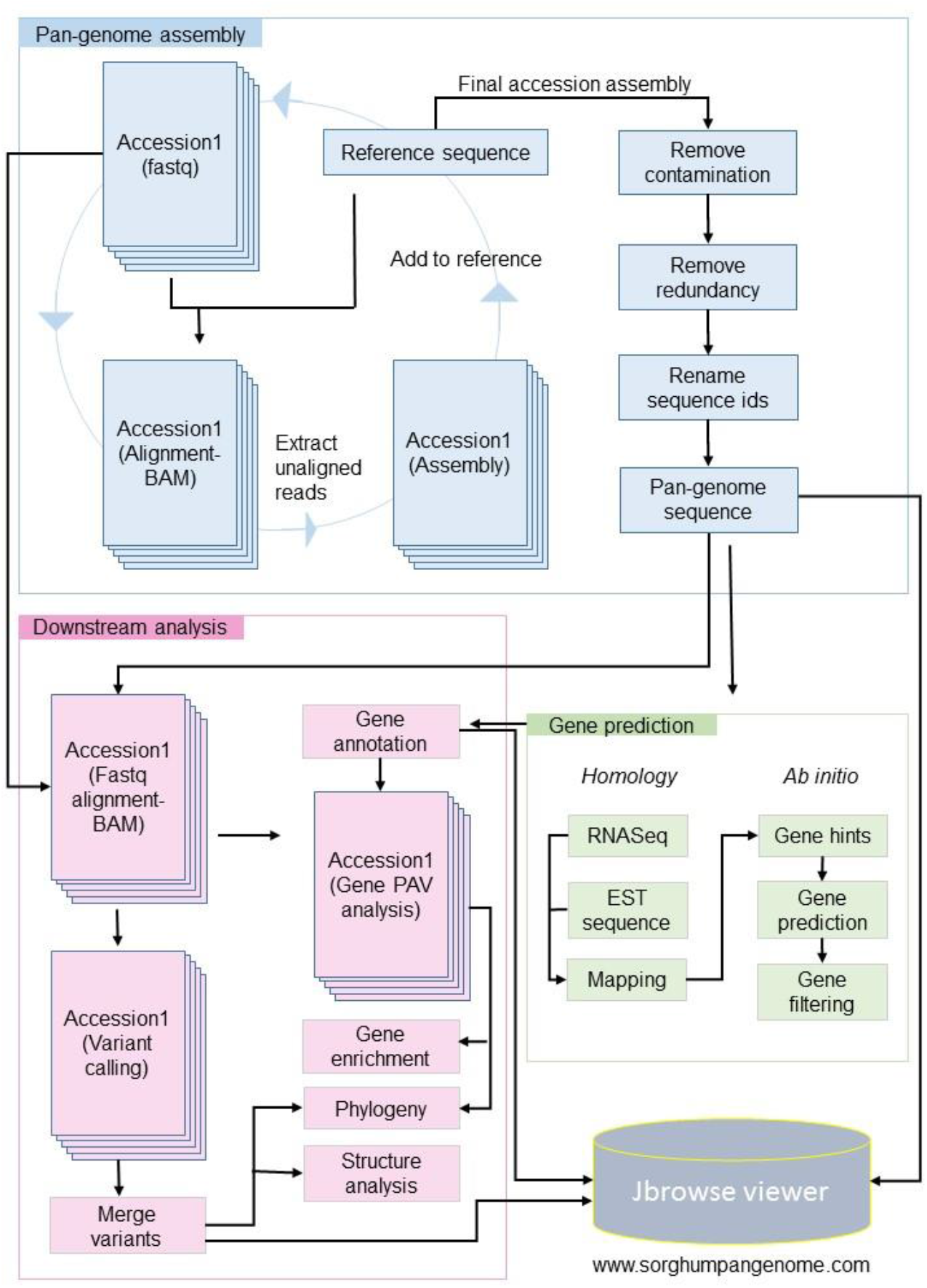
Raw reads of accession were aligned to reference, the unaligned reads were assembled and the contig sequence added to the reference sequence. The updated reference with assembled contigs used as reference for next iteration. The contamination and redundant sequence filtered from final updated reference and genes predicted with homology and *ab initio* method. On availability of pan-genome sequence and gene models, the downstream analysis performed including PAV, variant calling, phylogeny, enrichment and structure analysis.

On an average, each iteration of the process added 1.9Mb of sequence to the reference (Supplementary Figure 1) and a total of 263.7Mbp was assembled. Of these, 89.2Mb of the sequence were removed as contaminants (including chloroplast and mitochondrial sequences) and/or duplicated contigs. The final resulting pan-genome contained 210,805 contigs with a total length of 883.3Mb (Figure 2) with a minimum contig size of 500bp. Gene density on the contigs added by this pan-genome exercise was lower than on assembled chromosomes but comparable to the density observed on the reference unplaced scaffolds (Figure 3).

**Figure 2:**
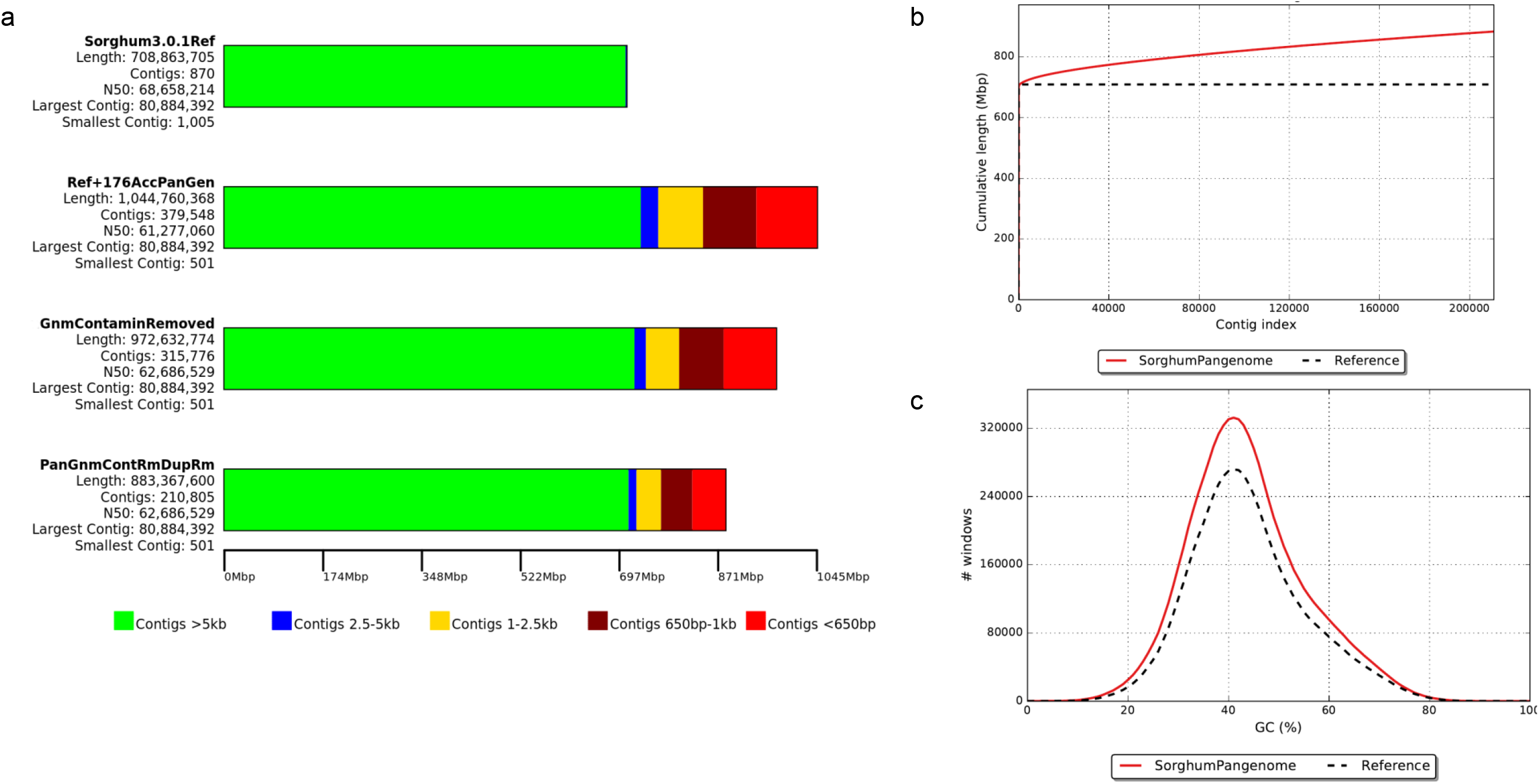
The development of sorghum pan-genome from reference genome assembly: **a)** a draft pan-genome (including sequence contamination, duplication, chloroplast and mitochondrial sequence) and final pan-genome assembly sequence size, b) cumulative length of assembled contigs and c) GC percentage of cleaned assembled contig sequences.

**Figure 3:**
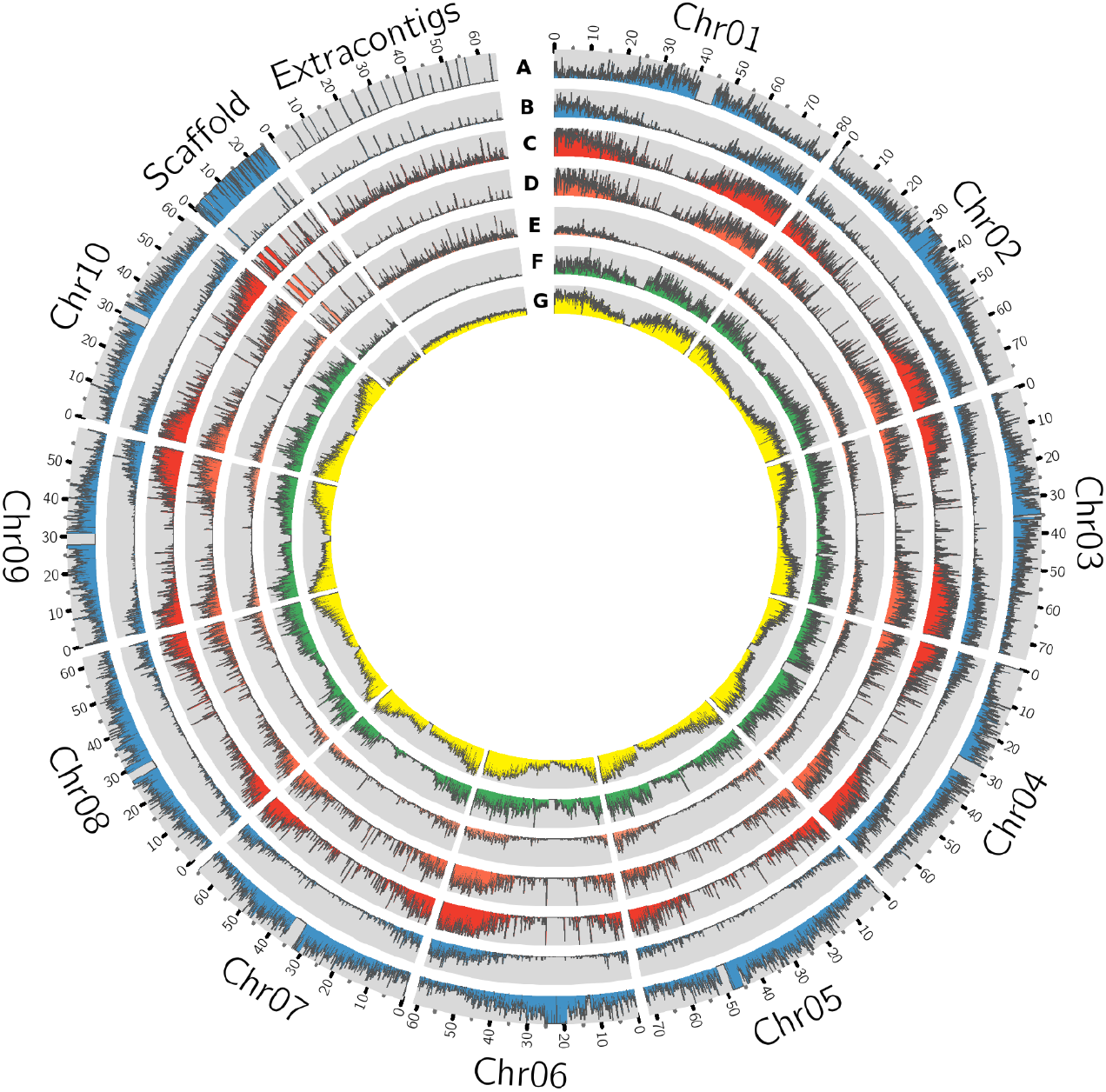
Circos plot of sorghum pan-genome having the extra contigs as a novel sequence assembled from 176 accessions. The genomic features of each track represent absolute values to the respective track (with 80Kb window size), **A**. Reference whole genome sequence reads mapping **B**. Drought expression (RNASeq) sequence mapping density. **C**. Gene density **D**. Genes commonly present in all accessions (core genes) **E**. genes absent in at least one of the accessions (variable genes) **F**. SNP density **G**. Insertions and deletions (indels).

The pan-genome showed an increase of 24.6% (174.5Mb) over the reference genome (Supplementary Table 2), which was the second-biggest increase of any previously reported pan-genome after the tomato pan-genome. The increase in tomato pan-genome was captured a 42% non-reference sequence from 725 accessions including the wild relatives (Gao *et al*., 2019). In other species, an increase in sequence size of 3.3% in wheat (Montenegro *et al*., 2017), 4% in *Oryza sativa japonica*, 6% in *Oryza sativa indica* (Yao *et al*., 2015), 5% in *Brachypodium distachyon* and 20% in *Brassica oleracea* (Golicz, Bayer, Barker, Edger, H. Kim, *et al*., 2016) was documented. The relatively large increase in sorghum pan-genome assembly size indicated that the presence of high level of genome diversity in the accessions.

The assembled sequence was annotated using a strategy called combining evidence-based *ab initio* gene prediction. RNASeq (Guo *et al*., 2019) mapping hints from the 25 accessions used for *ab initio* gene prediction and the 3,589 genes supporting the mapped expressed sequence tags (EST) sequences were retained. We identified 11,057 to 17,616 variable genes in the 176 genomes, with an average gene sequence length and exons per gene of 1,567bp and 3.6, respectively. The gene length and exons in core genes were more than the variable genes comparatively (Figure 4).

**Figure 4:**
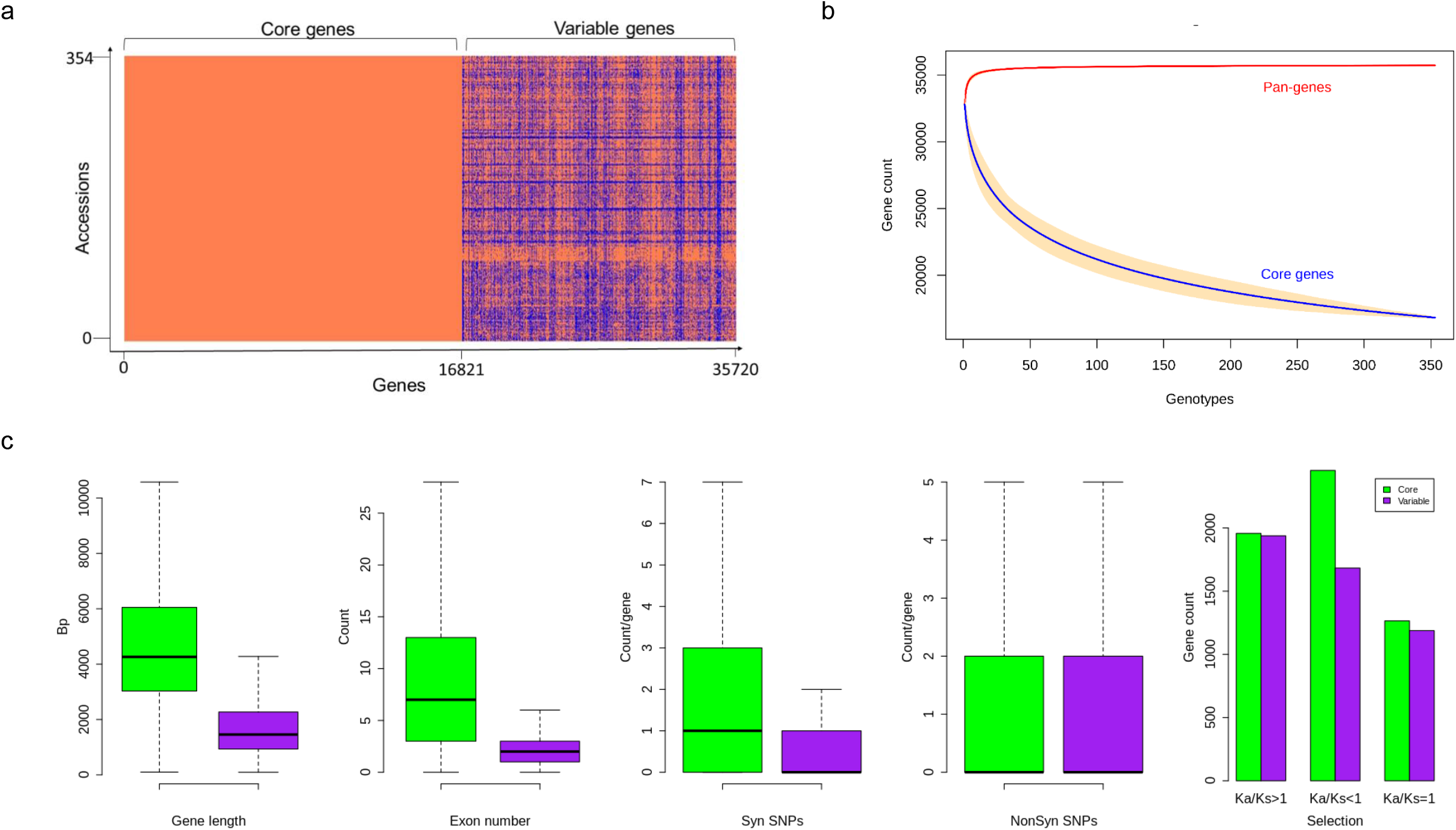
a) Gene presence and absence variations (gPAVs) in sorghum accessions b) Gene variation of pan-genome and core genome c) Sorghum core and variable gene properties. Variable gene length is shorter with fewer exons. The variable genes have fewer synonymous SNPs and similar non-synonymous SNPs compared to core genes. The gene counts with different Ka/Ks category indicating the selection pressure in both core and variable genes. There is a difference in the number of SNPs between core and variable genes within all groups.

### Sorghum pan-genome gene PAV (gPAV)

The gPAV in genes among the sorghum accessions could reveal the genetic changes that can be used to infer the phylogenetic history as well as to select the potential targets for breeding. To identify the gPAVs, sequence reads were mapped to the pan-genome contigs and genes were scored as present or absent based on the mapped sequence read coverage (Supplementary Table 3, Figure 5). For a given gene, to assess the gene loss event, the mapping of the whole genome sequence reads was measured. On an average, each sorghum accession contained 32,795 genes (Supplementary Table 4), of which 16,821 (47%) were core genes or in other words, they were shared by all remaining accessions. Comparatively, tomato (Gao et al., 2019) (74.2%), *Arabidopsis thaliana* (Contreras-Moreira *et al*., 2017) (70%), wheat (Montenegro *et al*., 2017) (64%), pigeon pea (Zhao *et al*., 2020) (86%) and, *Brassica napus* (Hurgobin *et al*., 2018) (62%) had higher number of genes (Bayer *et al*., 2020). On the other hand, 18,898 genes were variable/accessory genes (Figure 4.a), of which 30 genes were uniquely present and 3,183 (8.9%) were uniquely absent in any one of the accessions (Supplementary Table 3, Figure 5). Variable genes were found shorter and had fewer exons per gene when compared to core genes (Figure 4.c) which were in agreement with *O. sativa* and *A. thaliana* crop studies (Bush *et al*., 2014; Golicz, Batley, *et al*., 2016; Schatz *et al*., 2014). Based on gPAVs from 354 cultivars, we estimated the sorghum pan-genome had a closed type of pan-genome (Figure 4.b), with 30 genes were uniquely present and 3,183 genes were uniquely absent (Supplementary Table 5). The uniquely present genes were fewer than the wheat (49 unique genes per cultivar) (Montenegro *et al*., 2017) and *B. oleracea (*37 unique genes per cultivar) (Golicz, Bayer, Barker, Edger, H. Kim, *et al*., 2016). Of the 30 genes uniquely present in any single sorghum accession, nine such genes were reported from Macia accession alone (Figure 5). Extending the population size and including the wild relatives could further increase the measure of the gene content of this species (Figure 4.b) (Golicz, Batley, *et al*., 2016).

**Figure 5:**
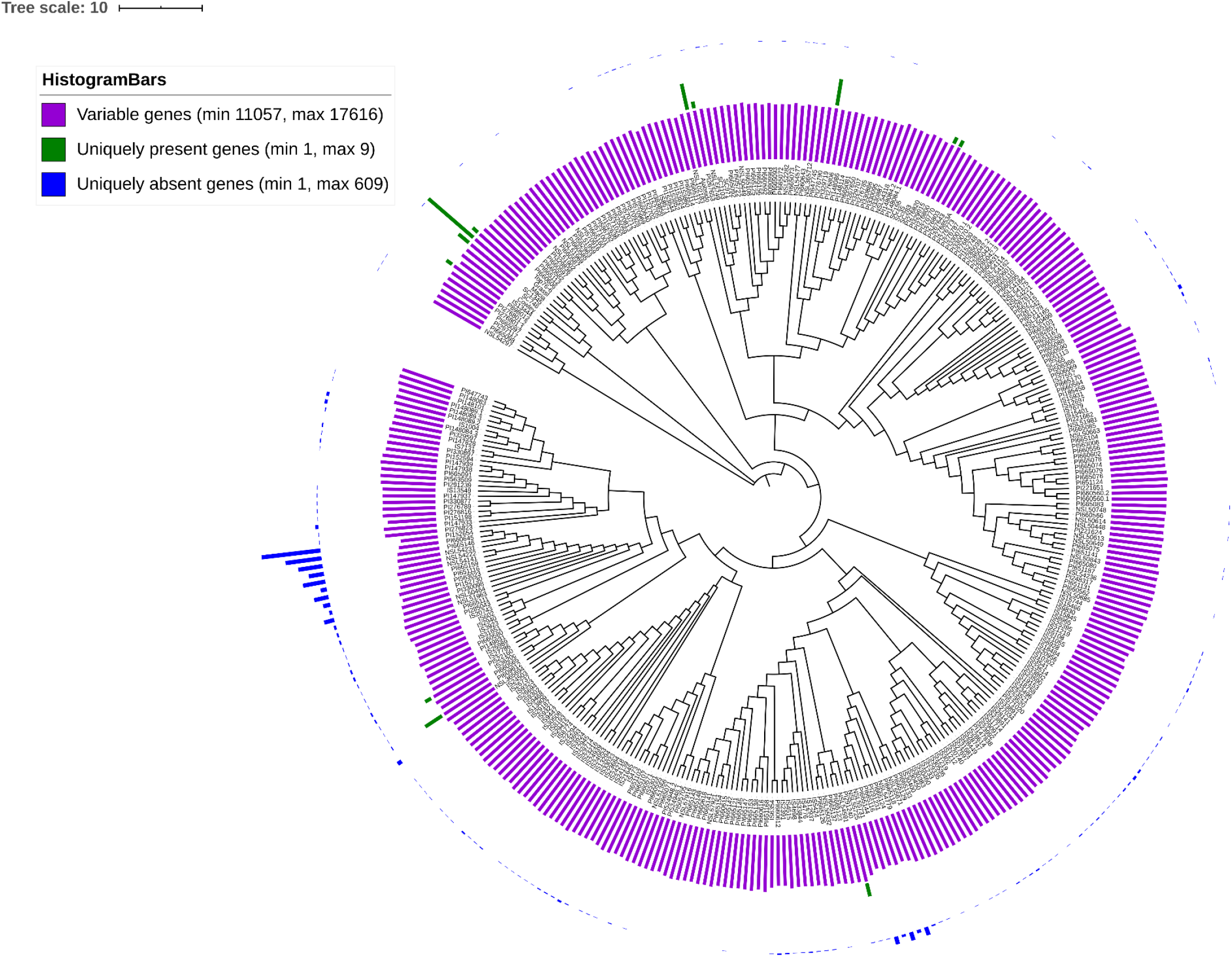
gPAVs-based neighbour-joining tree shows the genetic relationship among the sorghum accessions. Histogram shows the variable genes (purple bars), uniquely present genes (green bars) and uniquely absent genes (blue bars).

### Gene functional analysis

We identified enriched biological pathways by performing gene enrichment analysis using the R topGO package. The significantly enriched pathways related to responsive genes were identified (Figure 6). A total of 94 most significantly enriched genes (Supplementary Table 6) for biological process pathways are shown in Supplementary Figure 2. The gene ontology (GO) enrichment analysis showed that the genes were enriched in response to chemical, hormone, organic substance, stress, auxin and abiotic stimulus (Figure 6). It was noted that most of the pathways were related to plant response to stimulus and chemicals. The gene enrichment among stress responses genes including water deprivation (GO:0009414), desiccation (GO:0009269), abiotic stimulus (GO:0009628), chemical stimulus (G0:0042221) and stress (GO:0006950) were reported in a reference set of genes (Woldesemayat and Ntwasa, 2018). The gPAV-based enrichment on assembled genes from the sorghum pan-genome has added the response of the genes to auxin (GO:0009733), hormone (GO:0009725), organic substance (GO:0010033), hypoxia (GO:0001666) and decreased oxygen levels (GO:0036294). The functional annotation of the variable genes highlighted the genes involved in response to biotic and abiotic stress indicating the possible evolutionary adaptive traits (Lasky *et al*., 2015). Macia (9 genes), Ajabsido (4 genes) and PI329719 (4 genes) had a large number of unique genes (Figure 5), which could be used as potential donors for trait improvement. The functional analysis of such unique genes involved in response to the stimulus (GO:0050896), chemical (g8132, GO:0042221) and arsenic-containing substance (g24192, GO:0046685) (Supplementary Table 7).

**Figure 6:**
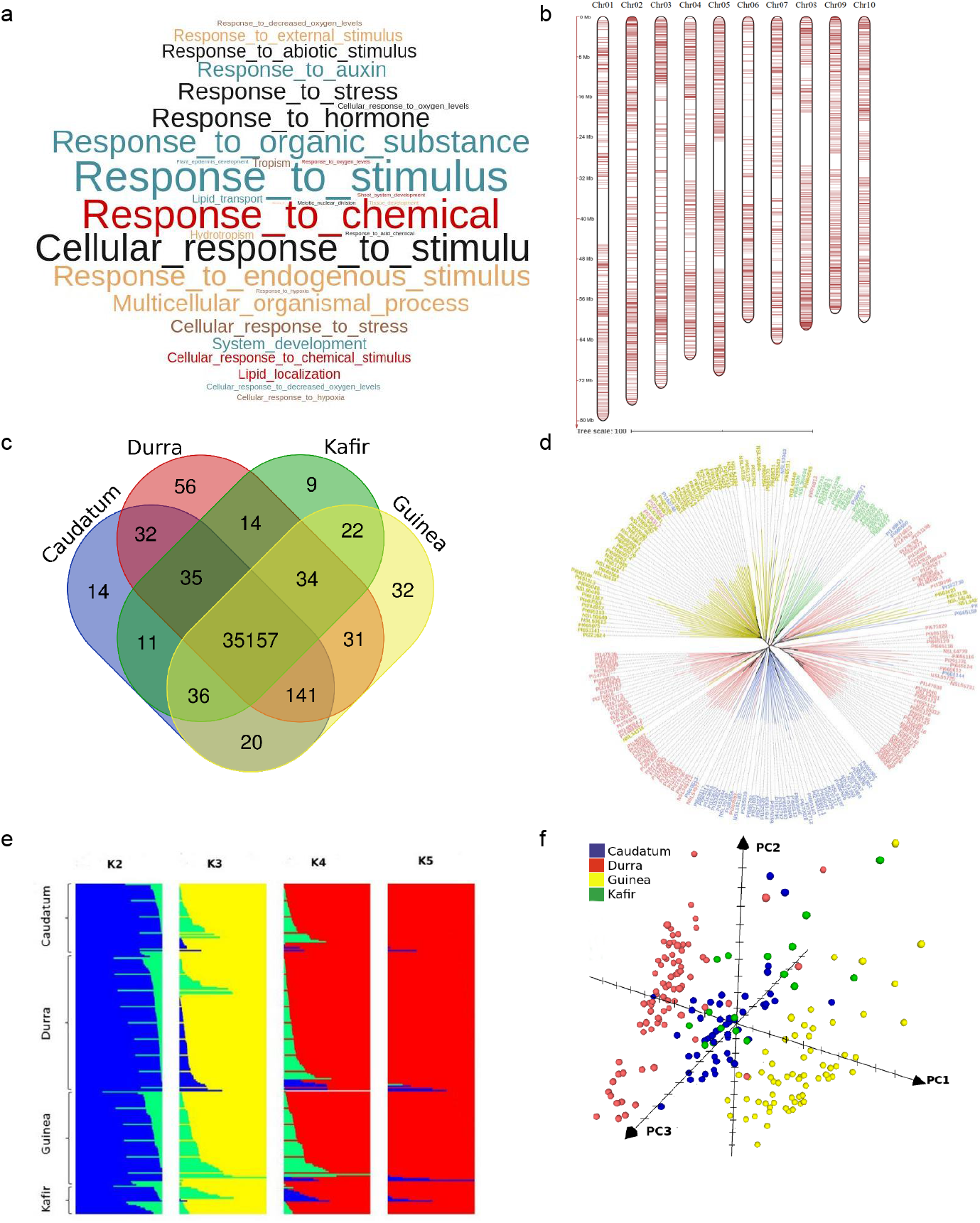
a) Significantly enriched top GO terms among the variable genes (p<0.05) b) distribution of Infinium SNP array markers on chromosomes c) specific and common genes across races d) neighbour-joining tree shows the genetic relation among the sorghum accessions belonged to different races (blue-*Caudatum*, red-*Durra*, green-*Kafir* and yellow-*Guinea*) e) structure analysis of sorghum population individuals with K2 to K5 and f) sorghum accessions grouped by *caudatum, durra, guinea* and *kafir* race through principal co-ordinate analysis (PCo).

**Figure 7:**
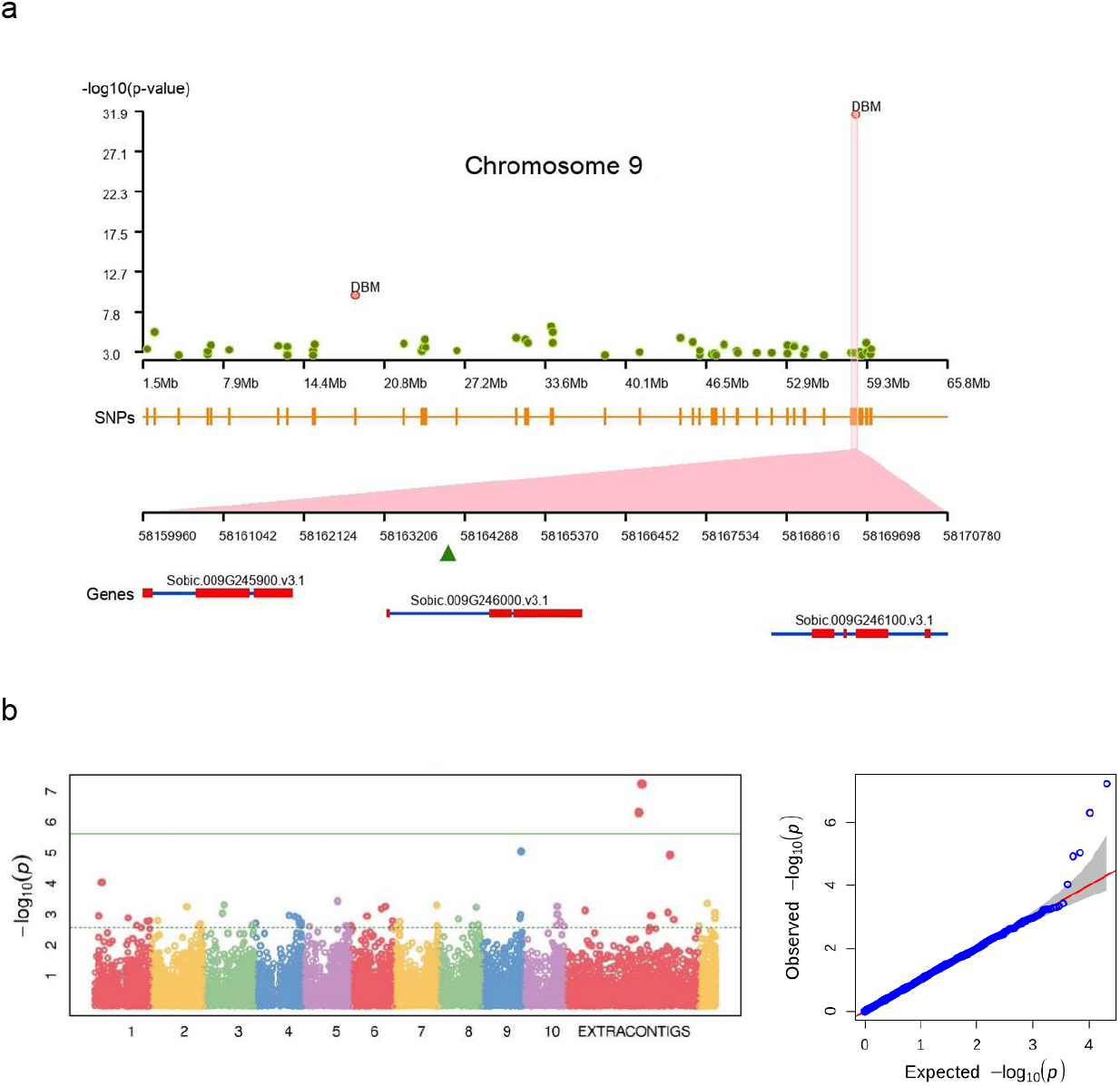
Genome-wide association showing the significant association of SNPs for **a)** plant biomass on chromosome 9 (on Sorbic.009G246000.v3.1) and **b)** plant height trait association on extra-contigs.

**Figure 8:**
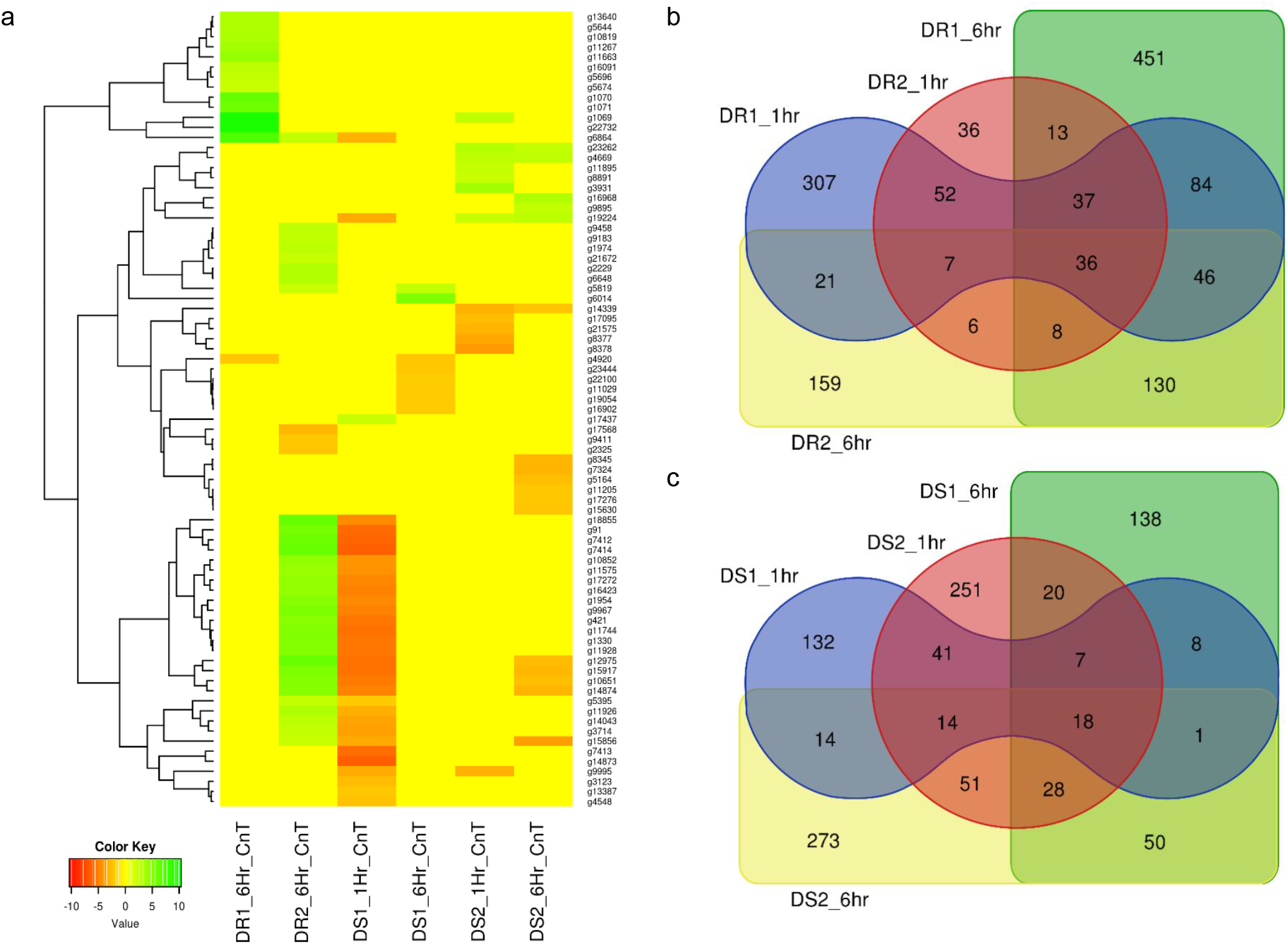
a)Sorghum drought trait-based RNASeq assay on pan-genome assembled genes (extra contigs). DR1 and DR2 were the two datasets for drought resistance after 1hr and 6hrs, respectively. DS1 and DS2 related to drought susceptibility after 1 hr and 6 hrs, respectively. Heatmap showing the range of up-regulated genes as green and down-regulated genes as red colour **b)** Drought resistance in BTx623 (DR1) and SC56 (DR2) and **c)** Tx7000 (DS1) and PI482662 (DS2) drought susceptible genes share between the genotype and treated conditions (1hr and 6hrs).

### Variant discovery

We identified a large number of variants (single nucleotide polymorphism (SNPs) and indels) by mapping the sorghum population whole-genome sequence reads to sorghum pan-genome assembly using GATK. Of the total of 2.0 million SNPs, 91,319 were in the extra contig (assembly) sequence (Supplementary Table 8). The SNP density in extra contigs (0.52/Kbp) was much less compared to the density in the reference genome assembly (2.72/Kbp) (Figure 3). The SNP annotation results illustrated the highest number of SNPs in intergenic region (40%) followed by upstream (22.5%), downstream (21.4%), intron (8.8%) and exon (3.6%) regions with an overall Ts/Tv ratio of 1.92. Chromosome 4 had the highest number of SNPs (251,830), followed by chromosomes 1, 2, 3, 5, 10, 8, 6, 9 and 7, respectively. Chromosome 7 had the fewest number of SNPs (119,019) with the highest density of 0.55/Kbp and chromosome 4 had the least density of 0.27/Kbp (Supplementary Table 8) (Figure 3). The SNPs and indels were highly dense in the telomere regions compared to centromeres explained the higher gene activity towards the telomeres. (Figure 3). The SNP annotation reported the frequency of synonymous SNPs in the core genes was much higher than in the variable genes (Figure 2). This was in contrast to the higher mis-sense SNPs in core pigeon pea genes to variable genes early reported (Zhao *et al*., 2020). We detected genome-wide indels of various size (Supplementary Figure 3.a) and the genes featuring indels has reduced proportionally to the size of the indels (Supplementary Figure 3.b). On increasing the indel size, the number of the indels decreases in both gene and genome-wide sequence. The overall indels count from the sorghum population used in this study was much higher than the indels earlier reported in the six sorghum lines (Yan *et al*., 2018). A total of 36,097 genes had 983,060 CNVs among the sorghum accessions used in this study. The Ka/Ks ratio estimating the balance between neutral mutations, purifying selection, and beneficial mutations on a set of core and variable genes exhibited that, core gene count under positive selection were significantly close to variable gene count compared to genes under the negative selection pressure (Figure 4).

The maximum (432,286) and minimum (2,854) number of SNPs were identified in sorghum accessions PI267614-NSL54318 and IS3693-IS23514, respectively (Supplementary Table 9). The accessions NSL54318 (849,052) and IS3693 (17,084) had the maximum and minimum polymorphism, respectively with sorghum pan-genome assembly sequence (Supplementary Table 10). The SNPs were validated with 3K SNPs Infinium array (Bekele *et al*., 2013) and of the 2,980 mapped flanking SNPs sequence, only 20 did not map on pan-genome assembly. The overall alignment rate was 99.33%, from the mapped 2,980 SNPs array sequences (Supplementary Table 11, Figure 6.b). Among them, 37 SNP sequences were mapped to extra contigs (novel sequence assembly) and 150 (5%) did not represent any GATK SNPs calls (29 SNPs from extra-contigs). In addition to the core SNPs of the array sequence, more SNPs were identified in the flanking sequence. Out of 15,383 GATK SNPs on the mapped array sequence, 15,314 SNPs were validated with the GATK called allele (Supplementary Table 11). Finally, on validation with array SNPs, the overall GATK SNP calling reported 99.9% accuracy.

To understand the genetic relationship of the 354 sorghum accessions, a neighbour joining (NJ) tree was constructed with the SNPs (Supplementary Figure 4). The accessions were arranged in many sub-groups indicating the possible sorghum race accessions. To assess the race-specific accessions, the 216 known sorghum race accession bootstrapped to construct an NJ tree. The NJ tree showed the subgroups of sorghum accessions according to the races with an exception of few individuals placed in other groups indicating the hybridisation process in the past (Figure 6.d). For example, PI221662, a durra race accession was genetically related to the *guinea* race. Similarly, to understand the gene PAV-based genetic relation, the phylogenetic relationship among sorghum accessions was assessed by distance-based 1000 bootstrap replicates and represented through the NJ tree. Among the 35,719 total genes, 53% exhibited the genic variations to estimate the relationship among the accessions (Figure 5). The largest number of genes uniquely present and absent genes was found in Macia (9 genes) and PI660645 (372 genes), which indicated the evolutionary distance from other accessions.

With the known four races (Obilana *et al*., 1996), the structure analysis with the variants set showed the presence of three sub-population (Figure 6.e), resulting an expected admixture between *caudataum* and *kafir* accessions which was in agreement with early study (Valluru *et al*., 2019) (Figure 4). The result was also validated by the PCo, where *durra* and *guinea* sorghum races displayed identifiable clusters, because of the available sequence representation through pan-genome, while *caudatum* and *kafir* individuals exhibited the admixers (Figure 6.f). The earlier principal component study shows the mixed grouping of *guinea* and *kafi*r accession in the (Sapkota *et al*., 2020), indicating the missing sequence representation for all race in the single reference genome. The group of *guinea* race accessions in PCo supported the Harlan and Stemler’s hypothesis, that the *guinea* race was probably the first diverged race than other races from early *bicolor* domestication events (The Races of Sorghum in Africa, 2012).

### Variation of sorghum race-variable genome

Sorghum pan-genome analysis has identified 18,898 variable genes, among which 111 were race-specific variable genes in the given population. The gene cluster analysis identified 11,470 gene families, of which un-clustered genes (6,057) included 556 from the non-reference genes and the remaining 5,501 were reference genes. Among these un-clustered genes, 3,195 were orthologous to *Zea mays, Setaria italica, Brachypodium distachyon*, and *Oryza sativa* and the remaining 2,862 were paralogous. The gene shares among four sorghum races showed that the *durra* and *guinea* had a maximum of 56 and 32 unique genes, respectively making them more diverse than the other two races with 14 (*caudatum*) and 9 (*kafir*) unique genes (Figure 6.c). The gene annotations suggested that the unique genes from *durra* were associated with heat shock protein, LRR repeat protein, L-type lactin-domain receptor, ABC transporter family proteins, and Ras-related proteins. *Guinea* group had the unique genes associated with disease resistance protein, beta-glucosidase proteins, NRT1/PTE protein family and Alpha/beta-Hydrolases superfamily proteins (Supplementary Table 12). The gene uniqueness to specific races possibly reflected the selection of the genotypes for adapting to the respective ecological conditions (Upadhyaya *et al*., 2017).

### Genome-wide association analysis (GWAS)

Two populations namely, Pop1 (Valluru *et al*., 2019) and Pop2 (Usha Kiranmayee *et al*., 2020) were used for GWAS to understand the functional utility of the pan-genome. Pop1 had 216 accessions with the phenotypes of dry biomass (DBM), plant height (PH) and starch (ST) while Pop2, a stay-green fine-mapping population with 190 segregates, had green leaf area (GLA), glossy (GL), plant vigour (V), leaf sheath pigment (LSP), shoot fly dead hearts (SFDH), trichome low (TL) and trichome up (TU).

In Pop1, the SNPs were further filtered by accessions and on applying the SNP quality filters, which retained 1.12 million SNPs for association analysis. Pop2 having sequence data of 190 genotypes processed to map to pan-genome and 109,338 SNPs were used for GWAS.

We identified a total of 397 unique SNPs having significant association (having p-value and false discovery rate below 0.05) in both Pop1 and Pop2 traits, of which 216 SNPs were commonly mapped with multiple traits. Most of these SNPs distributed on chromosome 10 (120 SNPs) followed by chromosome 6 (69 SNPs). The reference genome alone had 385 SNPs and the rest of the SNP-trait associations located on the unmapped read sequence assembly.

For the Pop1, a total of 36 SNPs had a significant association across three traits (Supplementary Table 13). Among them, seven were located on newly assembled contigs (DBM and PH), three were from unplaced reference contigs and the remaining 26 are from chromosome sequence. Among the 36 linked SNPs, 10 were genic and the remaining 26 were inter-genic regions (Supplementary Table 14). Three of the genic SNPs were associated with DBM while six were associated with PH and the remaining one co-mapped to both DBM and PH. From the 10 associated genes, three genes (Sobic.002G022500, Sobic.003G173400 and Sobic.004G350800) were from the core gene set and the remaining belonged to variable genes.

From the Pop2, the GLA, at various stages (Supplementary Table 15), associated with 219 SNPs including 111 genic, distributed across all chromosomes including the pan-genome assembly contigs (Supplementary Table 13). GL, LSP, SFDH, TL and TU traits were associated with 129 and 103 significant SNPs in *Rabi* (R13) and *Kharif* (K13) seasons, respectively (Supplementary Table 13). The majority of the SNPs were associated with chromosome 10 followed by 5 and 6 in both the seasons. Among them, a total of 96 SNPs was mapped across seasons and a total of 18 and 196 showed season-specific association in K13 and R13, respectively. Interestingly, only four genic SNPs were associated with TU in K13, whereas 63 were associated in R13 explained the season-specific gene regulations. Similarly, SFDH had no association in K13 but had 56 genic SNPs in R13 season (Supplementary Table 15).

The number of SNPs associated with DBM, PH, ST (Pop1), plant vigour (V), GL, LSP, SFDH, TL and TU, and GLA (Pop2) was 10, 25, 1, 1, 23, 31, 84, 169, 98 and 397, respectively. Among the chromosomes, as many as 392 of the SNPs were associated with chromosome 10 and only 8 SNPs were associated with chromosome 3 (3 SNPs on scaffolds). The pan-genome assembly contigs hold 15 trait-associated SNPs, an additional genetic resource for the sorghum breeding program.

Of the total 183 GWAS SNPs directly associated with gene functions, the DBM and PH (from Pop1) were associated with 10 genes (off these, 1 gene assembled in this study). In Pop2, 173 genes were distributed as 96, 11, 13, 46, 48 and 1 for GLA, GL, LSP, SFDH, TL and TU, respectively.

### Identification of the drought candidate genes

A sorghum RNASeq data generated from drought-resistant (BTx623 (DR1) & SC56 (DR2)) and susceptible (Tx7000 (DS1) and PI482662 (DS2)) genotypes at different seeding stages (Abdel-Ghany *et al*., 2020) were re-analysed and mapped through the newly developed pan-genome. A total of 1,788 genes were significantly affected by drought stress (Figure 3) and among them, 79 genes were reported from genes on assembly sequence (extra-contig) (Supplementary Table 17).

The drought-resistance (DR1 and DR2) and drought-susceptibility (DS1 & DS2) samples were phenotyped at two conditions (1hr and 6hrs). The DR1 and DR2 samples reported (1hrs treatment) a total of 590 (450 up and 140 down-regulated) and 195 (180 up- and 15 down-regulated) expressed genes respectively. Of these, none of the genes reported from the novel sequences, indicating both (DR1 and DR2) were closely related. Additionally, DR1 and the reference sequence belong to the same genotype and this supports the absence of gene expression from the novel sequence at this condition. When the treatment was extended for 6hrs, 14 (13 up and 1 down-regulated) and 34 (31 up and 3 down-regulated) genes from novel sequence were expressed for DR1 and DR2 data-sets respectively.

Similarly, DS1 and DS2 samples showed (for 1hrs treatment) 235 (123 up and 112 down-regulated) and 430 (388 up and 42 down-regulated) genes, respectively. Of these, DS1 and DS2 had 32 (1 up and 31 down-regulated) and 13 (7 up and 6 down-regulated) expressed genes from novel sequence, respectively. After 6 hrs of treatment, DS1 and DS2 samples reported 270 and 449 expressed genes respectively. Of these, DS1 and DS2 reported 8 (2 up and 6 down-regulated) and 17 (5 up and 12 down-regulated) expressed genes were from the novel sequence, respectively.

Over-all, five drought-related genes were co-mapped with the trait-associated genes. Among the five genes, three traits-linked genes Sobic.001G363200 (GLA), Sobic.007G180300 (GL) and Sobic.010G231900 (TL, TU and SFDH) were commonly expressed in drought resistance and susceptibility conditions. The remaining two drought resistance specific genes Sobic.005G069800 and Sobic.006G127800 were linked to PH and LSP traits.

## Discussion

We built a sorghum pan-genome with an iterative mapping and assembly approaches with 176 of 354 whole genomes sequenced accessions having coverage of more than 10X. The total size of the pan-genome has become 883Mbp, with a 20% increase (175Mbp) compare to the reference assembly of 708Mbp. This level of novel sequence increase probably due to the high level of genetic diversity observed in the respective species (Cuevas and Prom, 2020).

We have generated the pan-genome genomic resource from the diverse sorghum accessions including the basic and intermediate sorghum races (bicolor (B), *caudatum* (C), *durra* (D), *kafir* (K) and *guinea* (G)). Comparison of the wide range of sorghum whole genome sequence datasets has enabled to assemble many coding genes that were absent in published sorghum reference genome sequences. The mapping of RNASeq read from 25 accessions on the assembled contigs supports the predicted genes on the novel sequence (Supplementary Figure 5) is an additional genetic resource that will enhance the identification of the QTL and genome-wide association studies (Chen *et al*., 2014),(Yano *et al*., 2016),(Zhao *et al*., 2020). The earlier pan-genome studies found that non-reference genes have significantly involved in agronomic traits mainly in plant defence responses (Dolatabadian *et al*., 2020; Golicz, Bayer, Barker, Edger, H. Kim, *et al*., 2016; Hirsch *et al*., 2014; Montenegro *et al*., 2017). Similar to the sorghum genes, *B. oleracea* pan-genome genes also showed that nearly 30 percent of reference genes exhibited the gPAV (Golicz, Bayer, Barker, Edger, H. Kim, *et al*., 2016). It is understood that, as the number of genotypes increases, the size of core genes decreases with a relative increase of variable genes (Figure 4.b). With the 10 sample sizes, *Brassica oleracea* pan-genome had 20% of PAV genes (Golicz, Bayer, Barker, Edger, H. Kim, *et al*., 2016) which was in consistent with simulation with similar population size in *B. distachyon* pan-genome (Gordon *et al*., 2017). Similarly, pan-genome from 15 Medicago genomes had 42% of sequences share with few accessions (Zhou *et al*., 2017), which was comparatively similar to the 49% of sorghum variable pan-genes in this study.

The result of the structure groupings correlated with the PCo showing three different clusters with one of them having two groups (*caudatum* and *kafir*). Among the four basic sorghum races used in this study, PCo displayed, *guinea* and *durra* remain as distinct clusters while *caudatum* with *kafir* classified with mixed genotypes, which is considered as the stable hybrid race among the 10 possible stable combinations of sorghum races (Obilana *et al*., 1996). Similarly, mixed PCo clusters were also reported earlier with five basic sorghum races, where the sorghum B race was not well supported genetically and a majority of them share membership with the remaining four genetic groups (Brown *et al*., 2011).

The genomic features helped the races to group into different clusters. By looking at the race-specific genomic data, each race had distinctive features. The *guinea* group had 37 accessions with race specific genes present in range of 2 to 13 genes per accessions, whereas *durra* had 2 to 12 genes in 92 accessions, *kafir* had 2 to 5 genes in 12 accession and *caudatum* had 2 to 4 genes in 15 accessions. The two groups including *durra* and *guinea* were having 56 and 32 distinct genes, respectively unique to these groups, whereas *caudatum* and *kafir* have on 14 and 9 distinct unique genes, as these groups have the admixture accessions which share genes between the groups.

The functional analysis of variable genes was enriched with GO terms associated with response to light, flower development, salt stress, water, heat, desiccation, temperature, osmotic stress, lipid, gibberellin, and stress. The results supports the earlier gene function based clustering and enrichment analysis exhibiting the similar stress respons genes reported in sorghum (Woldesemayat and Ntwasa, 2018). The plant hypersensitive response annotation in the variable gene was reported in plant pan-genome analysis (Golicz, Bayer, Barker, Edger, H. R. Kim, *et al*., 2016; Hurgobin *et al*., 2018; Montenegro *et al*., 2017; Zhao *et al*., 2020).

The development and application of sorghum SNPs have limited to reference genome assembly sequence used in the analysis. The 1.8 million SNP reported earlier on Rio with respect to BTx623 (Cooper *et al*., 2019), were limited to the single reference genome. Using the whole genome sequence data from 354 sorghum diverse accessions, we identified two million SNPs and 3.9 million indel sites, which represented the functional genome diversity. The density of genetic variation in the novel assembled sequence was low compared to the reference sequence. The reference genome carried most of the conserved essential genes, indicating that the variable sequence has low diversity (Figure 3), as reported in the six sorghum landrace individuals from common geographical regions (Yan *et al*., 2018). The fewer number of SNPs on variable sequence mainly contained genes involved in response to various stress (biotic and abiotic stress tolerance), this is in consistent with the SNPs from disease resistance R genes differentiating sweet and grain sorghum accessions (Zheng *et al*., 2011). A reference sequence within the pan-genome assembly alone accounted for 95.4% of SNPs and the added assembly sequence from the sorghum population had 4.5% additional SNPs. A total of 2,980 array SNPs from (Bekele *et al*., 2013) were identified as similar to GATK called (reference-based variant calling) SNPs with 99.33% of true SNPs. The GATK called sorghum SNPs validation rate with array SNPs was higher (99.33%) compared to the non-reference based variant calling methods, for example, the wheat pan-genome SNPs were called with 96.3% accuracy (Montenegro *et al*., 2017). The abundance of SNPs depends on factors such as mutation events and genome diversity and the SNPs identified in the variable genome can assist in characterizing novel metabolic pathways.

Phylogenetic analysis of 354 sorghum accessions using SNPs on the pan-genome demonstrated the mixed groups of diverse biomass genotypes (Valluru *et al*., 2019), domesticated accessions (Guo *et al*., 2019) and Chibas sorghum breeding program accessions (Jensen *et al*., 2020). gPAV-based phylogeny showed a group of 15 accessions having uniquely absent genes in a range of 2 to 509 genes from the biomass genotypes indicating the wider genetic diversity. The five Chibas sorghum breeding lines (Macia, Ajabsido, SC1345, P898012 and Grassl) had the most unique genes followed by seven domesticated lines distributed across the phylogenetic tree. On assessing the known sorghum race genotypes from (Valluru, et al. 2019) phylogeny showed a cluster for each sorghum race. Few individuals of *caudatum* and *guinea* were mixed with *durra* cluster indicating that these are the *caudatum*-*durra* (CD) and *guinea*-*durra* (GD) hybrid individuals. Similarly, few accessions were not placed in respective race groups, for example, PI248317 accession was a *durra* race accession placed in *guinea* race cluster which shared the genetic similarity with *guinea* race as DG hybrid individual.

The GWAS performed in the earlier study was limited to the phenotype association only with limited SNPs on the reference genome used (Kimani *et al*., 2020; Morris *et al*., 2013). The SNP calling on sorghum pan-genome has enabled to identify the variants also from non-reference sequence assembly from the genetically diverse accessions. A total of 91,339 SNPs reported from the assembled sequence were the additional markers used for GWAS. A total of 36 SNPs (from Pop1) were associated with target traits. Among them, 10, 25 and 1 were from assembly sequence (extra contigs) were associated with DBM, PH and ST, respectively. Additionally, the GLA (from Pop2) had a significant association with five SNPs on extra-contigs. The GLA phenotypes in 2013 and 2014 after 7, 14, 21, 28, 35, 42, 49 days after flowering (DAF) in *rabi* were associated with 219 SNPs. Most of the SNPs were linked with the GLA recorded at the early stage of 7 (linked with 150 SNPs) to 14 DAF (linked with 161 SNPs) (Supplementary Table 13). From the flowering stage to 14 days of post-flowering, the GLA expression was significantly linked with 101 common SNPs (two SNPs reported from Extra-Contig101123 at 855 positions and Extra-Contig170379 at 501 base position) (Supplementary Table 13). For the phenotypes of GL, LSP, SFDH, TL and TU in both rabi and kharif seasons, a total of 147 SNPs were identified, of which 85 were co-mapped in both seasons, 44 were unique to *rabi* and 18 were unique to *kharif*. Out of total 397 associated SNPs, 12 SNPs from novel sequences having significant trait association is an additional gain from the pan-genome assembly.

Most of the associated SNPs linked to genes including NAC-domain protein controls the flowering time and stress response, BTB domain for protein-protein interaction, PSII protein complex for oxygenic photosynthesis, AAI domain protein for lipid transfer protein (LTP). The genes are transcription factors (TFs) such as nuclear TF, reverse transcriptase Ty1/Copia-type domain and BZIP. The genes also associated with ubiquitination pathway proteins such as B-box, F-box, U-box, RING-type and RING-type E3 ubiquitin transferase protein supporting the sorghumFDB gene family classifications (Tian *et al*., 2016).

We found 1,788 drought-responsive genes with different seeding stage sequence data mapping on pan-genome assembly, whereas weelky sampled the growing plants and mapping the RNASeq data to reference alone reported the 44% of genes exhibiting the response to drought stress (Varoquaux *et al*., 2019).

This difference in drought expression was expected between the seedling samples (in 1 to 6hrs difference) compared to root and leaf large scale sampling in 2 to 17 weeks of pre and post-flowering drought responses (Varoquaux *et al*., 2019). Similar drought stress gene expression changes were seen in laboratory and greenhouse studies in sorghum genotypes (Fracasso *et al*., 2016; Johnson *et al*., 2014). Identifying 79 drought-linked differentially expressed genes on assembly sequence are the additional genes added from this study (Supplementary Table 18). These additional genes through pan-genome were mainly involved in the cell membrane, catalytic activity, molecular function regulation, response to the stimulus, metabolic process, cellular and biological regulation.

## Methods

### Pan-genome assembly and annotation

The pan-genome was assembled using iterative mapping and assembly approaches. The assembly was initiated with a sorghum reference assembly v3.0.1 to map sorghum accession whole-genome sequence data iteratively. Reads from 176 sorghum accession were mapped to the sorghum reference v3.0 (McCormick *et al*., 2018) using Bowtie2 (Langmead and Salzberg, 2012) v2.3.4, and unmapped reads were assembled with IDBA_UD assembler (Peng *et al*., 2012) and the assembled contig sequence more than 500bp length was only considered and appended to reference genome sequence. The resulting final assembly sequence was compared with NCBI non-redundant nucleotide databases using BLASTn and the sequences with homology to sorghum mitochondria (NC_008360), chloroplast (MK348612) also the sequences having homology outside the green plant group Viridiplantae taxonomy group (Taxonomy ID: 33090) were removed. The remaining sequences were self-compared with nucmer search (http://mummer.sourceforge.net/) and sequences with greater than 90 percent coverage with greater 90 percent identity were removed to maintain the non-redundancy of the novel sequences. REPEATMASKER-v4.0.7 (Smit, AFA, Hubley, R & Green) masked repetitive elements using sorghum as the species. The sorghum expressed sequence tags (ESTs) from GenBank were aligned with tBLASTx and genes were predicted using AUGUSTUS v3.3.2, supporting the EST alignments. The gene models having fewer than 300bp in length were filtered out and the remaining genes supporting either EST alignments or hisat2 (Kim *et al*., 2019) alignments (RNASeq read from 25 accessions, Supplementary Table 1) further used for functional annotation against uniref90 (database downloaded in May 2020).

### Gene presence-absence variations (gPAVs)

Whole-genome sequence reads of 354 sorghum lines were mapped with Bowtie2 v2.3.4 (Langmead and Salzberg, 2012) to pan-genome assembly with a wide insert size range between 0 and 1000bp. The gene PAVs were defined based on sequence reads coverage mapped to respective genes as described by (Golicz, Bayer, Barker, Edger, H. Kim, *et al*., 2016). Genes models on contigs longer than 1Kbp were used in this analysis. PAV converted into binary matrix and with 1000 bootstrap resampling were used to estimate the genetic relationship among the accessions with R “ape” package (Paradis *et al*., 2004) to construct a NJ tree and visualized in iTOL tree viewer (Letunic and Bork, 2019).

The core genes were defined as the genes present in all the accessions, whereas the variable genes are the genes missing in one or more accessions. The *in-house* developed script was used to define the core and variable genes from the PAV matrix. Core and variable genes were compared for gene length, exon number, synonymous SNPs, non-synonymous SNPs and Ka/Ks. The mean count for each sample size of core and pan-genes present in all possible combinations of 354 accessions was plotted. The protein sequences of *Zea mays, Setaria italica, Brachypodium distachyon*, and *Oryza sativa* were downloaded from the public database UniProt for cluster analysis. All protein sequences were compared using all-by-all BLASTp followed by MCL for gene clustering into gene families with default parameters. The gene enrichment analysis was performed with Fisher exact test from R “topGO” package (Alexa *et al*., 2006) using “Elim” method.

### SNP discovery and annotation

The sorghum whole genome sequence reads of 354 accessions were quality trimmed using Trimmomatic (Bolger *et al*., 2014) and mapped to pan-genome using Bowtie2 v2.3.4 (Langmead and Salzberg, 2012) allowing to map paired reads. The aligned reads in SAM format converted to BAM format using samtools (Li *et al*., 2009) followed by filtering out the read duplication with Picard tools (http://broadinstitute.github.io/picard).Variants against the reference (pan-genome) were called with GATK v.4.1 (McKenna *et al*., 2010) and directed to quality filtered with vcftools v.0.1.13 (Danecek *et al*., 2011). The variant sites having missing genotypes of more than 0.15 and minor allele count less than two were excluded and the remaining sites were used for downstream analysis such as SNP functionally annotated with SnpEff v.4.3 (Cingolani *et al*., 2012).

### Sorghum diversity and population structure

A panel of 216 diverse sorghum accessions, including the four races, were used for genetic diversity and population structure assessment. A total of 1.12 million filtered SNPs from sorghum race accessions were retained for downstream analysis. The STRUCTURE v2.3 (Hubisz *et al*., 2009), was used to estimate the population structure using admixture model. The tested K was set from 2 to 5 and optimal K for population structure was defined with the structure program. With the same SNP set, PCo analysis was done with R labdsv package (https://CRAN.R-project.org/package=labdsv) and phylogeny analysis performed using 1000 replicates with R “ape” package (Paradis *et al*., 2004) and visualized in iTOL tree viewer (Letunic and Bork, 2019).

### GWAS

Two different mapping populations having the phenotypic data of ten traits were used for the association study.

#### Pop1

The phenotype and genotype data associated with PH, DBM and ST were adapted from (Valluru *et al*., 2019) for GWAS analysis. Among the 354 WGS data used for the above analysis, 227 genotypes belonged to four major races of sorghum having a representation from Africa, Asia and America. The SNPs corresponding to the above-mentioned genotypes were filtered with vcftools and used for GWAS. In 2016, the PH was recorded from 4 to 16 weeks after planting with an interval of 2 weeks, DBM and ST was measured at harvest.

#### Pop2

The stay-green fine-mapping population developed by crossing an introgression line cross RSG04008-6 × J2614-11 (Usha Kiranmayee *et al*., 2020) was used for association study using the pan-genome assembly. The DNA from parents and 152 individuals were isolated and skim-sequenced to produce genotype data to a depth of 0.1X. The sequence reads were QC’d with trimmomatic (Bolger *et al*., 2014), mapped with bowtie2 (Langmead and Salzberg, 2012) and SNP called with GATK (McKenna *et al*., 2010) and filtered with vcftools (Danecek *et al*., 2011) as above said method.

The Pop2 was evaluated with GLA trait in the *rabi* season of 2012-2013 and 2013-2014 at ICRISAT, Patancheru, India. The GLA percentage was measured from seven to 49 DAF for every seven days interval in both years. Additionally, in the year 2013, the phenotypes of GL, LSP, V, TL, TU, SFDH traits were recorded in rabi (R13) and kharif (K13) seasons.

The genotype to phenotype association was performed with GAPIT (Lipka *et al*., 2012) and the results were initially filtered with Bonferroni cut-off (-log10(p-value)>2.5) followed by *p*-value and false discovery rate values less than 0.05 (close to Benjamini-Hochberg cut-off value) as the significant values. On parsing the gene coordinates, for each candidate SNP within the range of genes were identified and assigned to SNP.

### Drought RNASeq assay analysis

The RNASeq data derived from two genotypes of drought-resistant (BTx623 (DR1) & SC56 (DR2)) and drought susceptible (Tx7000 (DS1) and PI482662 (DS2)) at the seedling stage was obtained from (Abdel-Ghany *et al*., 2020) SRA database. The quality check was performed on raw sequence reads using FastQC (Andrews, 2015) followed by cleaning the low-quality reads and removing sequencing adaptors using the Trimmomatic (Bolger *et al*., 2014) tool. Trimmed reads were aligned to the Sorghum pan-genome using TopHat2 (Kim *et al*., 2013) and bam files were filtered to remove reads aligned to multiple locations. Differential gene expression was performed on different conditions using Cuffdiff (Trapnell *et al*., 2010) to compute logFC and q-values across all lines at different conditions (control & treated). A total of eight conditions were analysed to find drought-induced genes after 1hr and 6hrs of post-treatment (20% PEG (polyethyleneglycol) treatment). Two biological replicates were analysed for each condition resulting in 32 samples (4 genotypes × 2 conditions × 2-time points × 2 replicates). The differentially expressed genes (DEGs) were determined if the q-value is less than 0.05 and log2FC is <-2 or >2 ratios between control and treatment for each time point and in each genotype.

## Conclusions

We constructed and characterised the sorghum pan-genome using the reference genome assembly and the whole-genome sequence reads of genetically diverse sorghum accessions. The pan-genome had 35,719 predicted genes, which were categorized as core, conserved genes, and variable genes as they exhibited presence and absence variation. The variable genes were enriched with genes response to various stresses. The SNP Infinium array result showed 99% of representation on the pan-genome assembly sequence. About two million SNPs were developed through pan-genome which can use for functional downstream research. The pan-genome resources were validated by assessing the genetic diversity of sorghum races, identification of genes from GWAS and RNASeq studies. These newly generated genomic resources could be used in sorghum genetic gain improvement programs.

## Acknowledgments

The authors thank Plant Breeding and Genetics Section, School of Integrative Plant Science, Cornell University, Ithaca, New York, USA and Plant Genome Mapping Laboratory, University of Georgia, Athens, Georgia, USA for providing the sorghum WGRS, RNASeq data and phenotype data. The authors also acknowledge the supporting funds from AVISA, ICAR and BMGF.

## Author contributions

PR and AR conceived and designed the project. PR, PG and SS carried out the analysis. SS managed computational resources and data management. SDP provided sorghum trait data. PR, NT, DE, GM, NB, RG and AR jointly wrote the manuscript. All authors have seen and the manuscript hasn’t been published elsewhere.

---

**Supplementary Figure 1:**
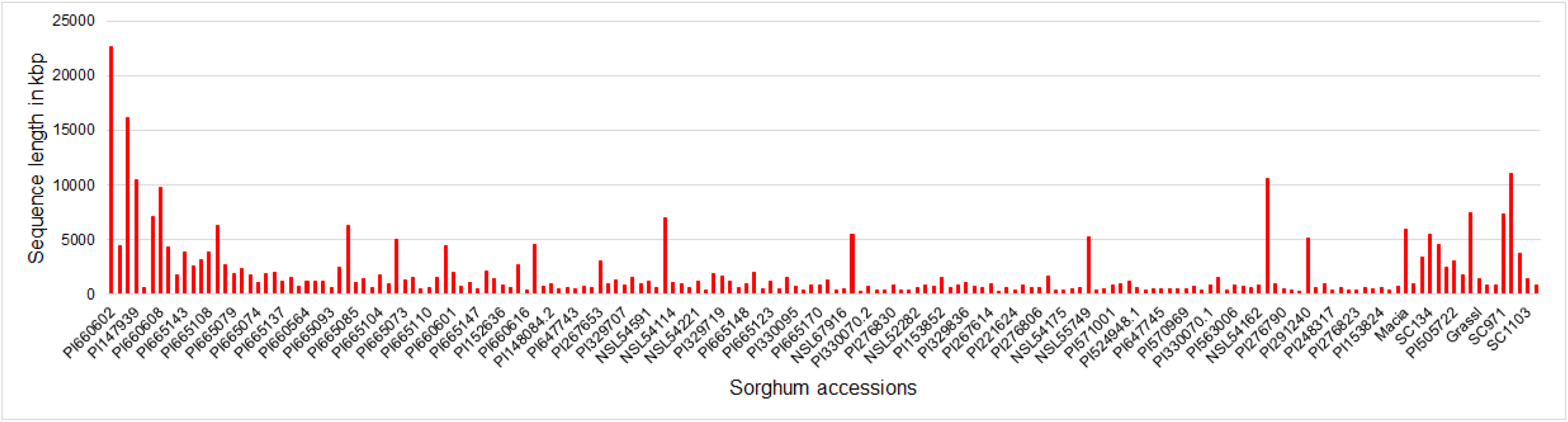
Whole genome sequence of 176 sorghum accessions mapped iteratively to the updated reference sequence assembly and the unmapped sequence reads were assembly iteratively. The plot represents the size of the sequence assembly gained from respective accessions.

**Supplementary Figure 2:**
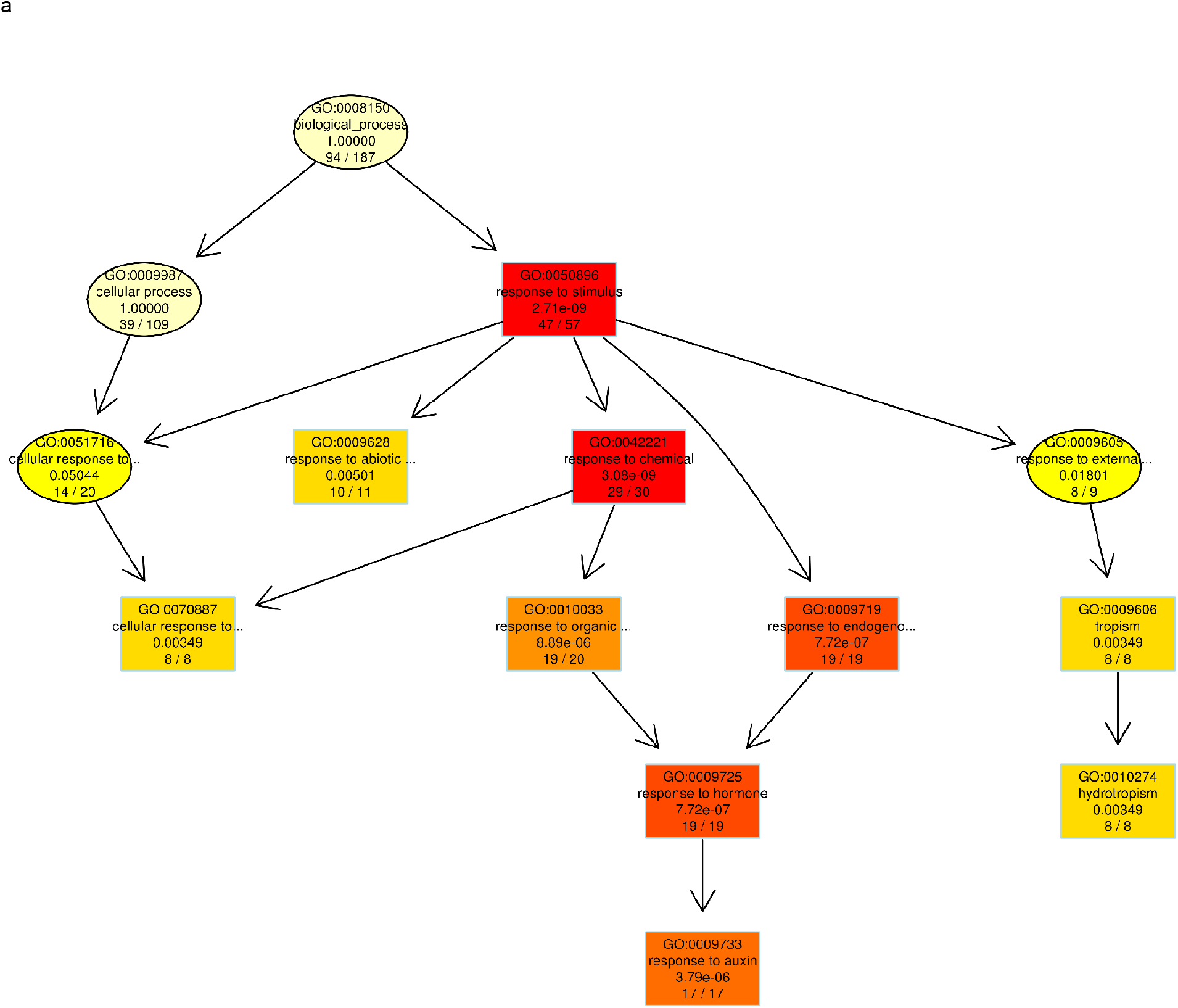
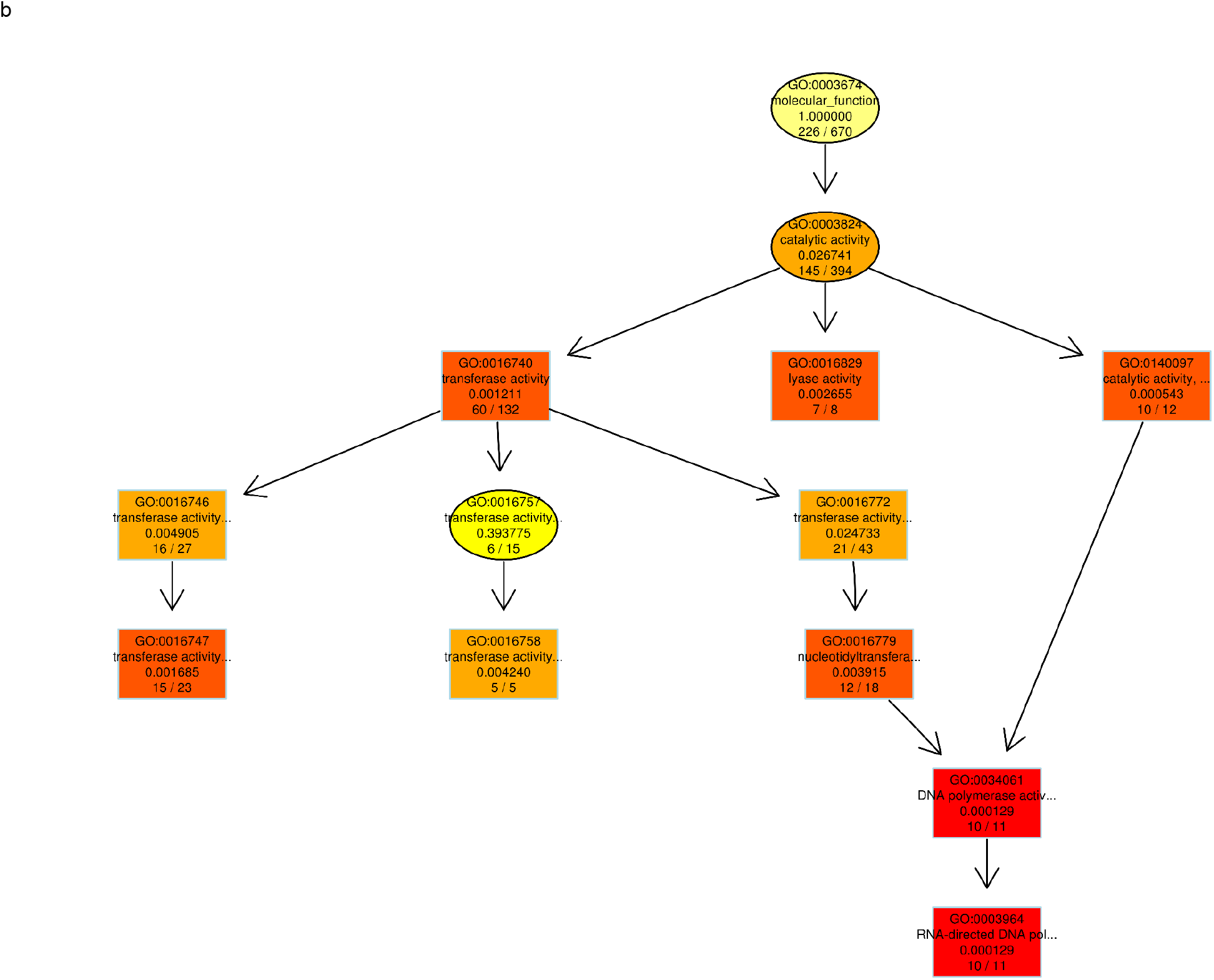
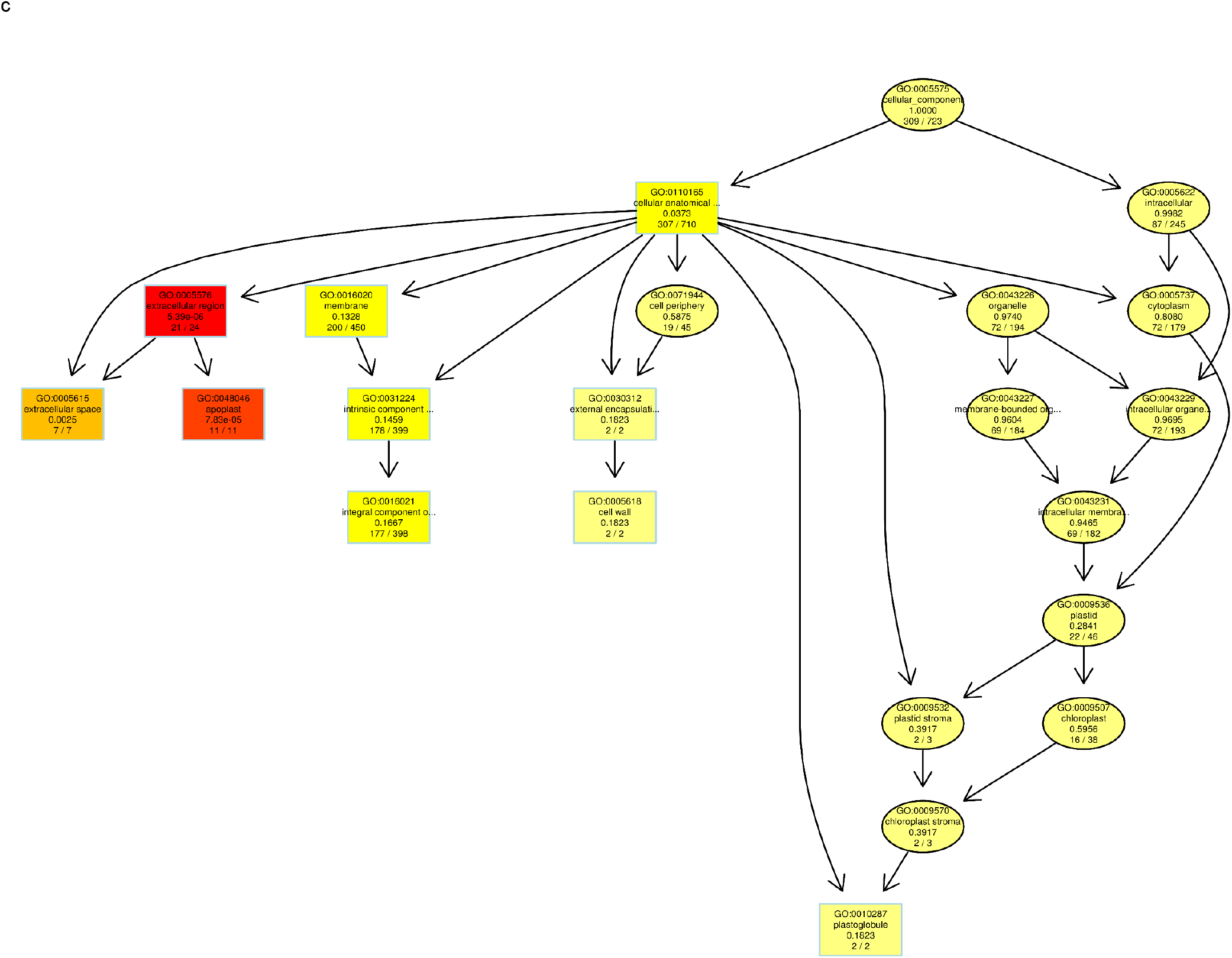
The sorghum pangenome variable genes enrichment analysis and the metabolic pathways for a) Biological process b) Molecular function and c) Cellular components. Top 10 GO terms identified for scoring GO terms for enrichment. Rectangles indicate the 10 most significant terms and the colour represents the relative significance ranging from red (most significant) to yellow (least significant). Each node has GO identifier and name with raw p-values and number of significant genes out of total genes annotated.

**Supplementary Figure 3:**
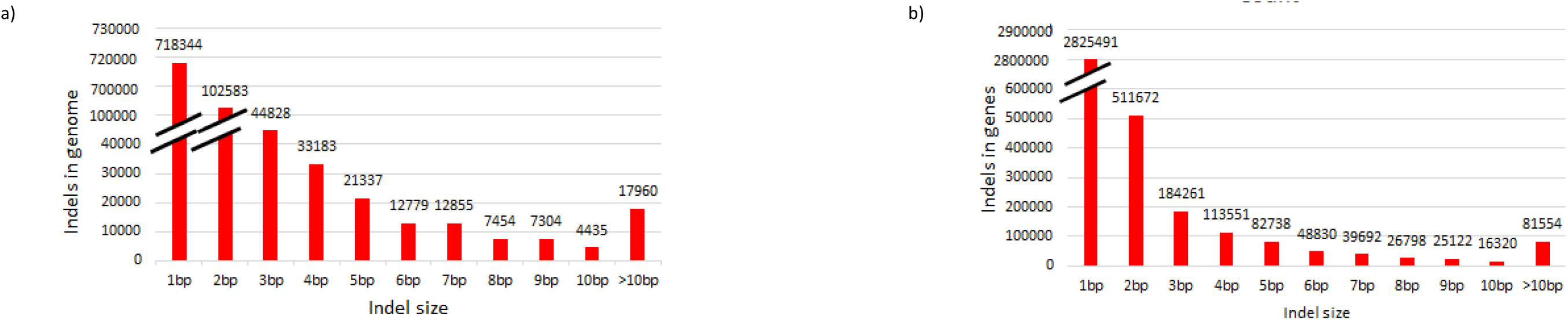
Insertions and deletions of various size distribution at sorghum pangenome a) genome level and b) gene level

**Supplementary Figure 4:**
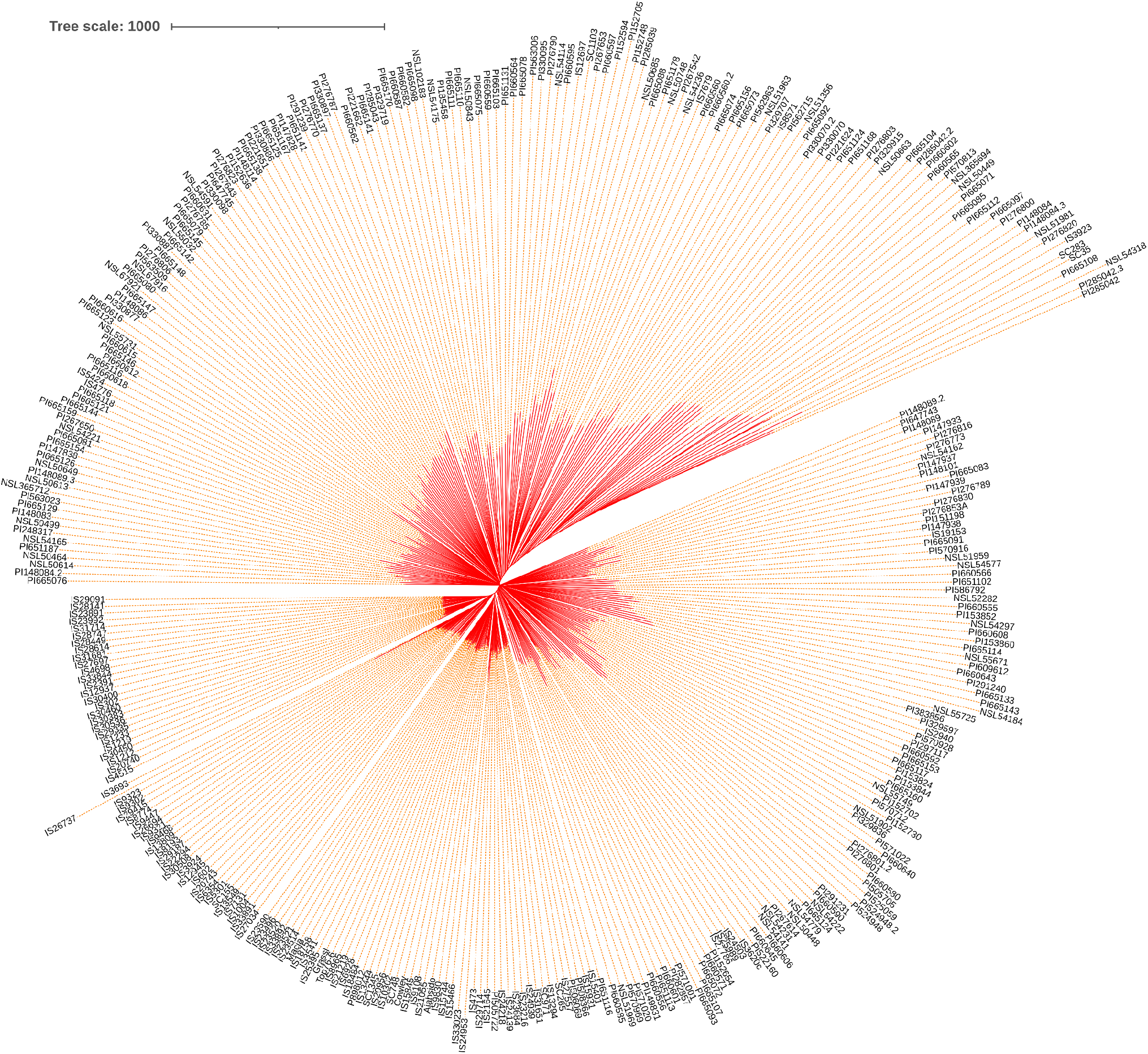
The genetic relationship of 354 sorghum accession neighbour joining (with 1000 bootstrap) analysis (the unrooted tree with branch length in red colour).

**Supplementary Figure 5:**
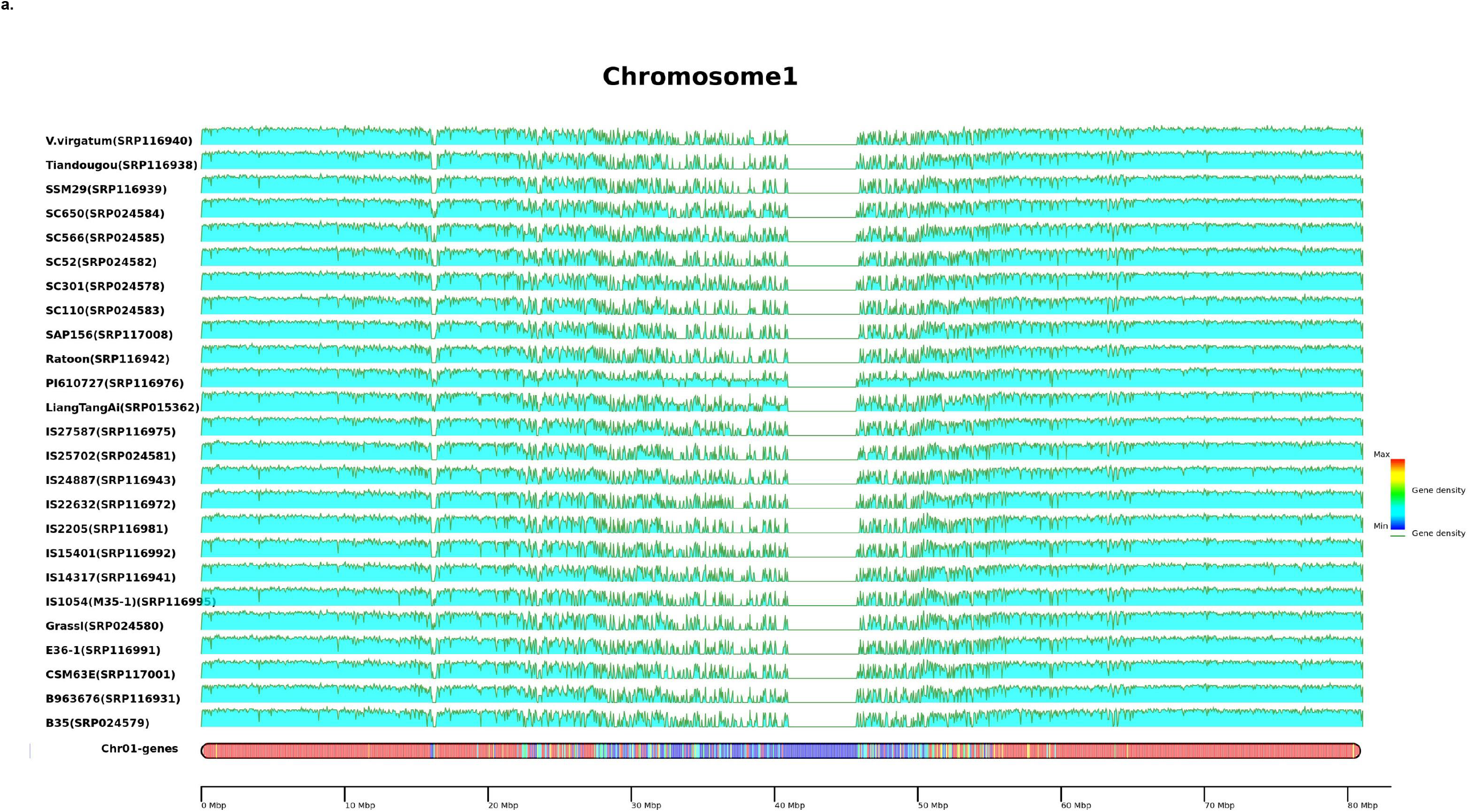
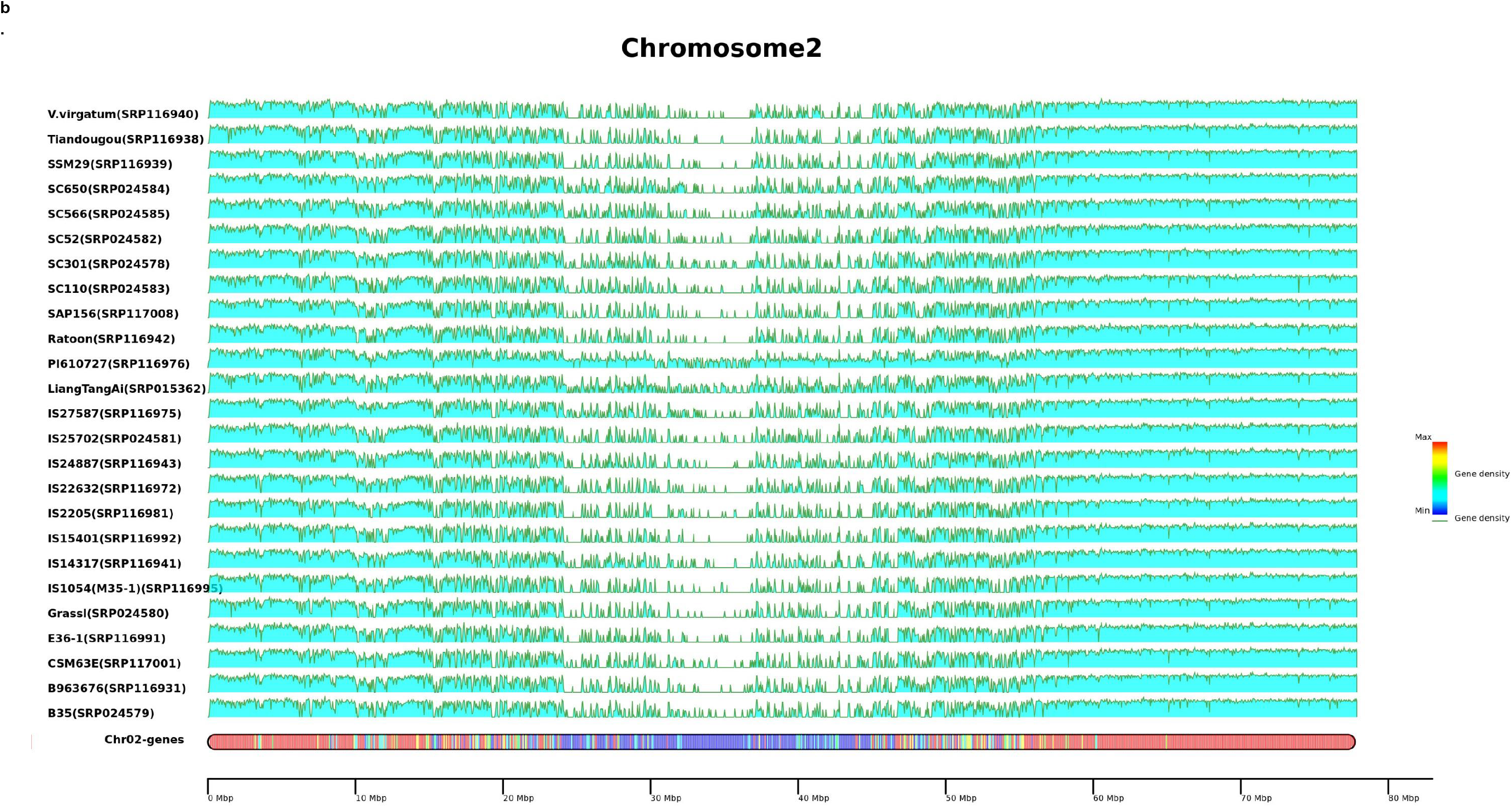
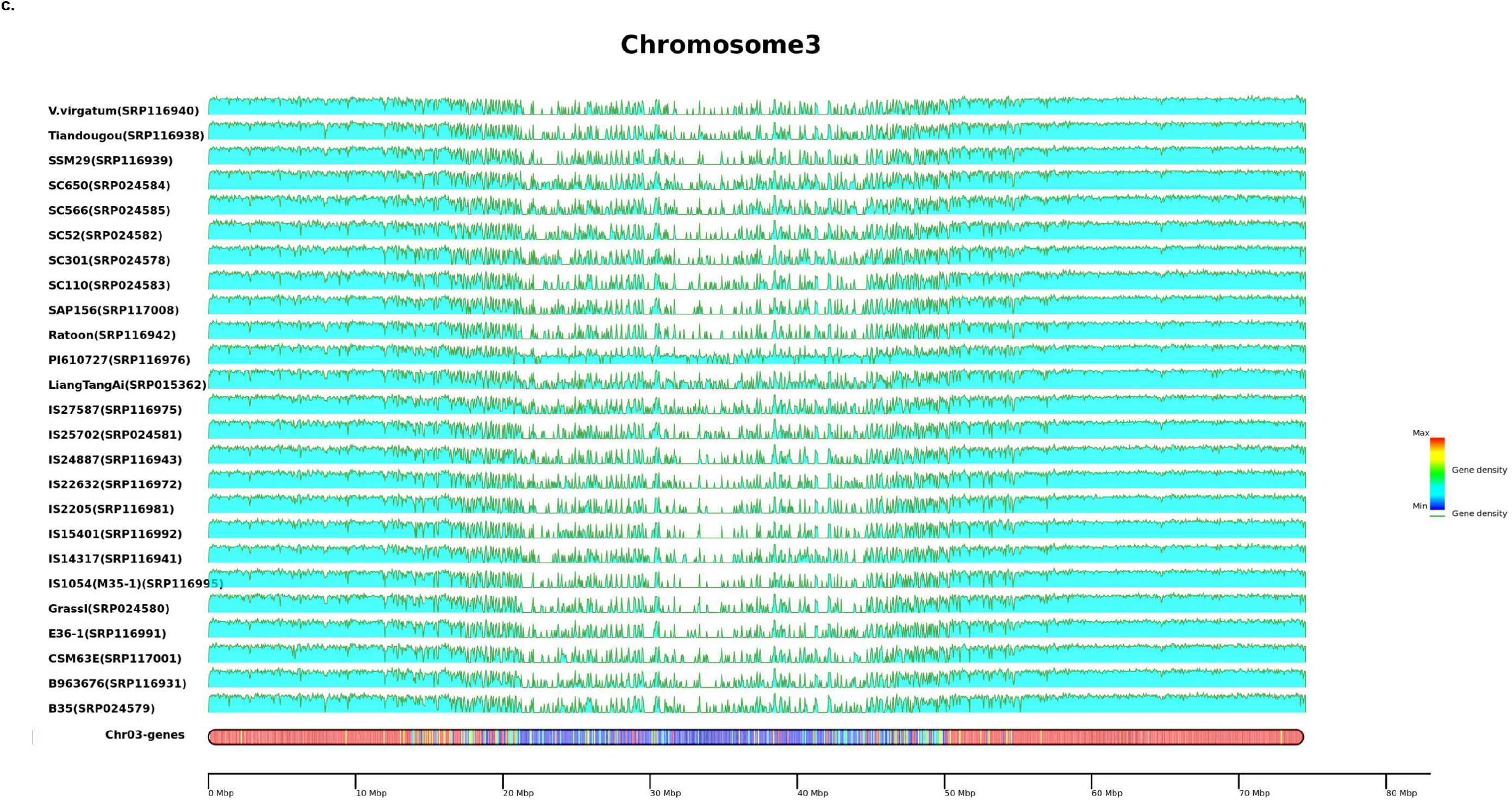
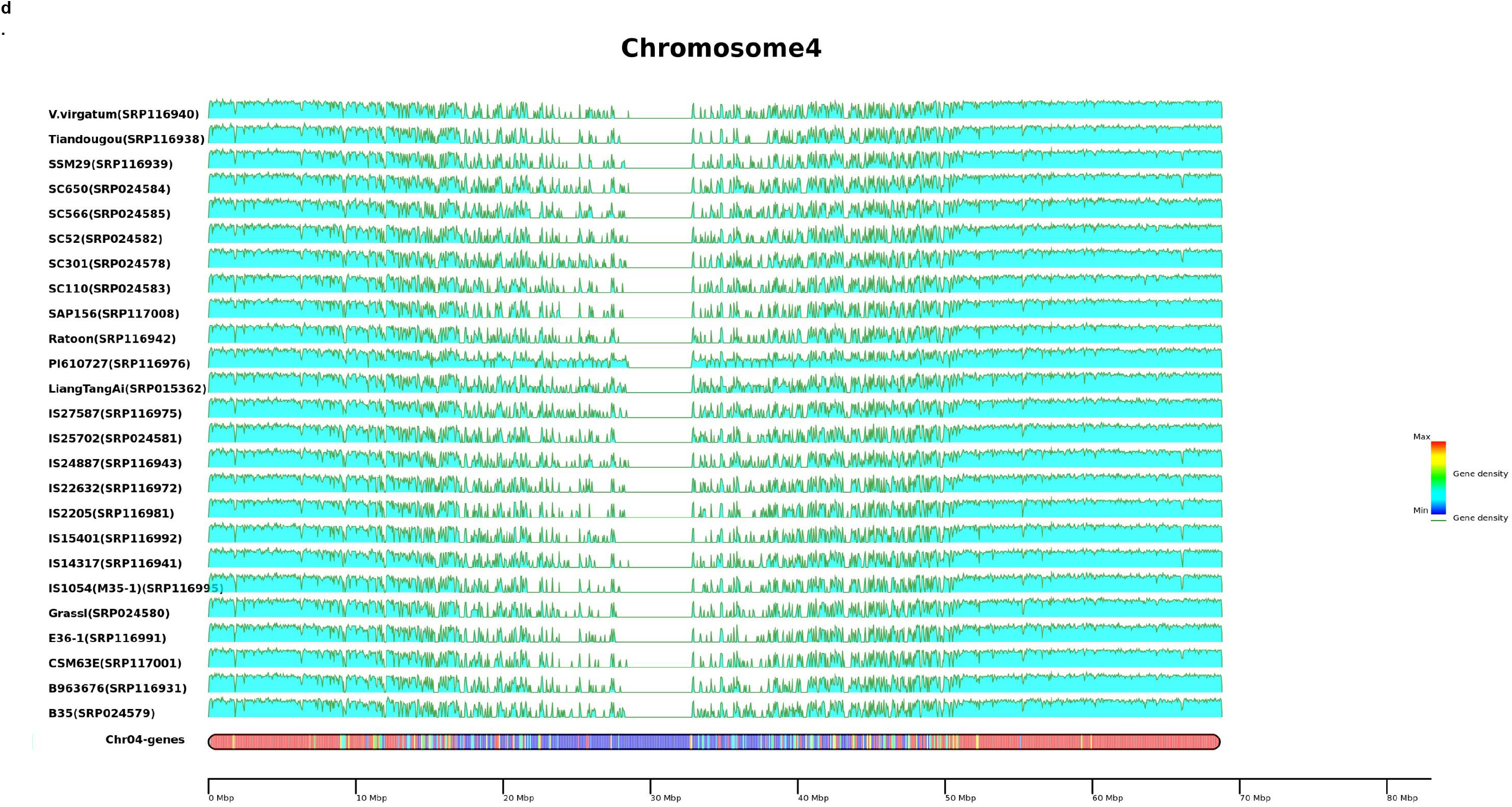
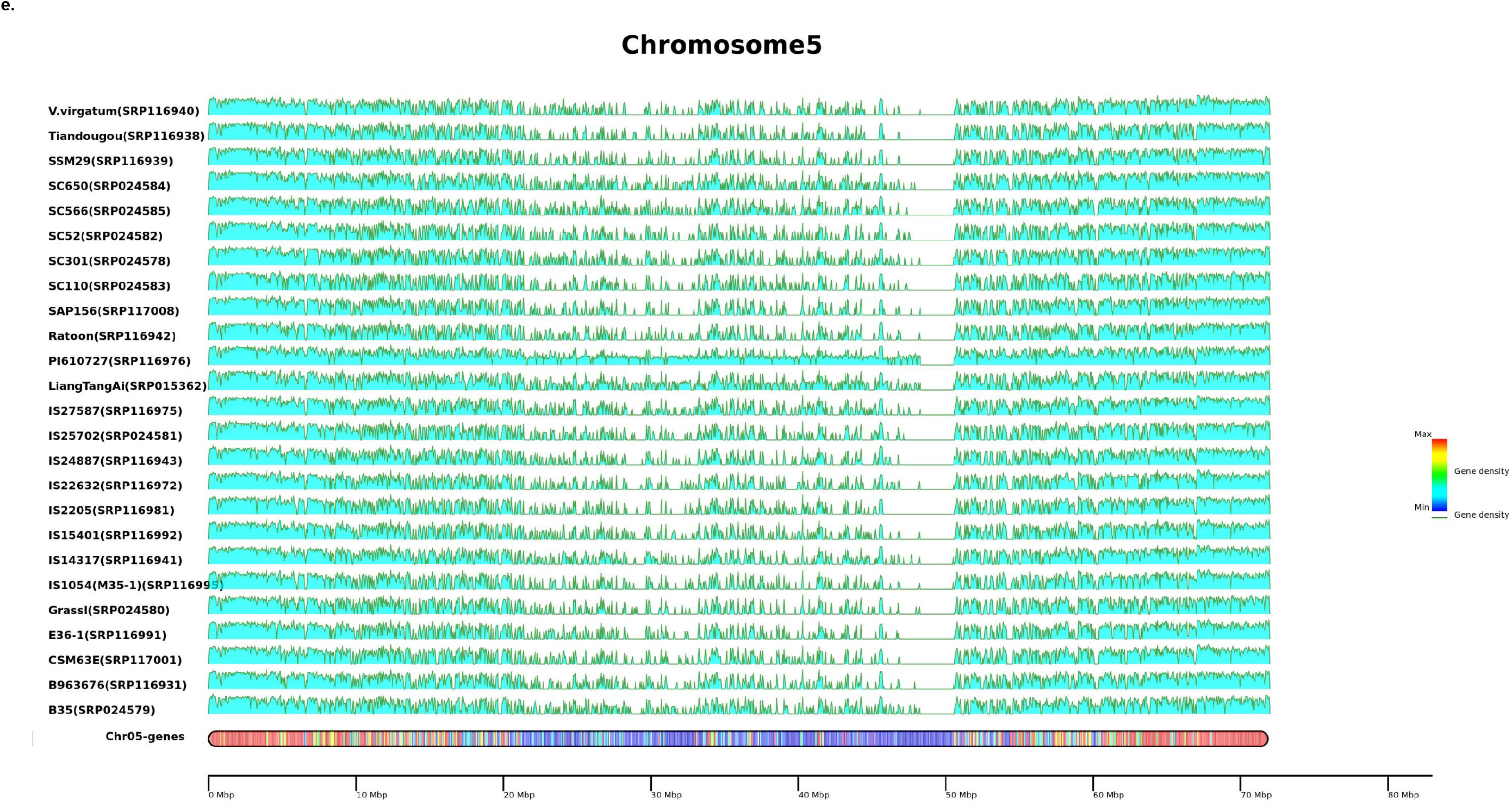
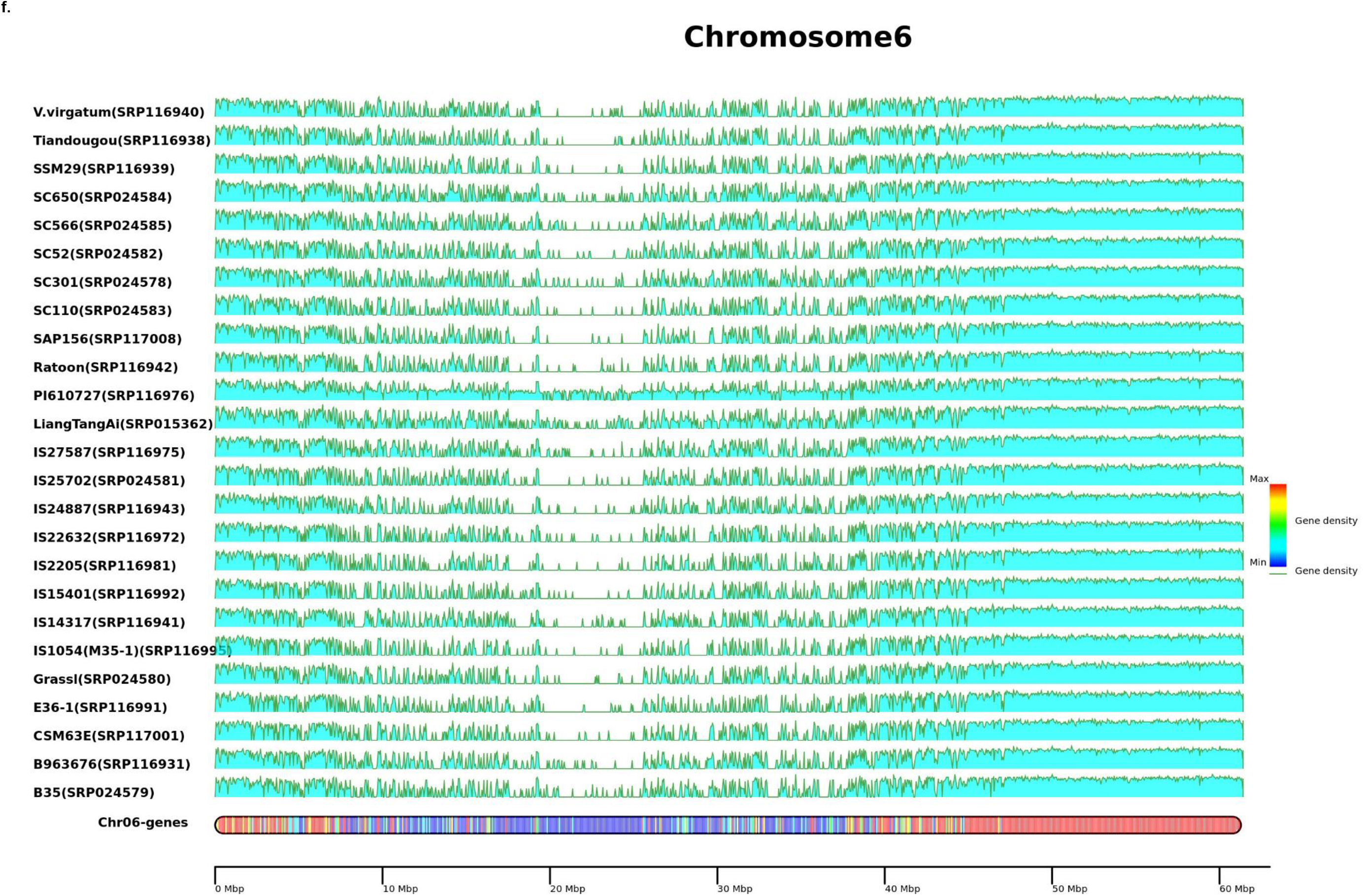
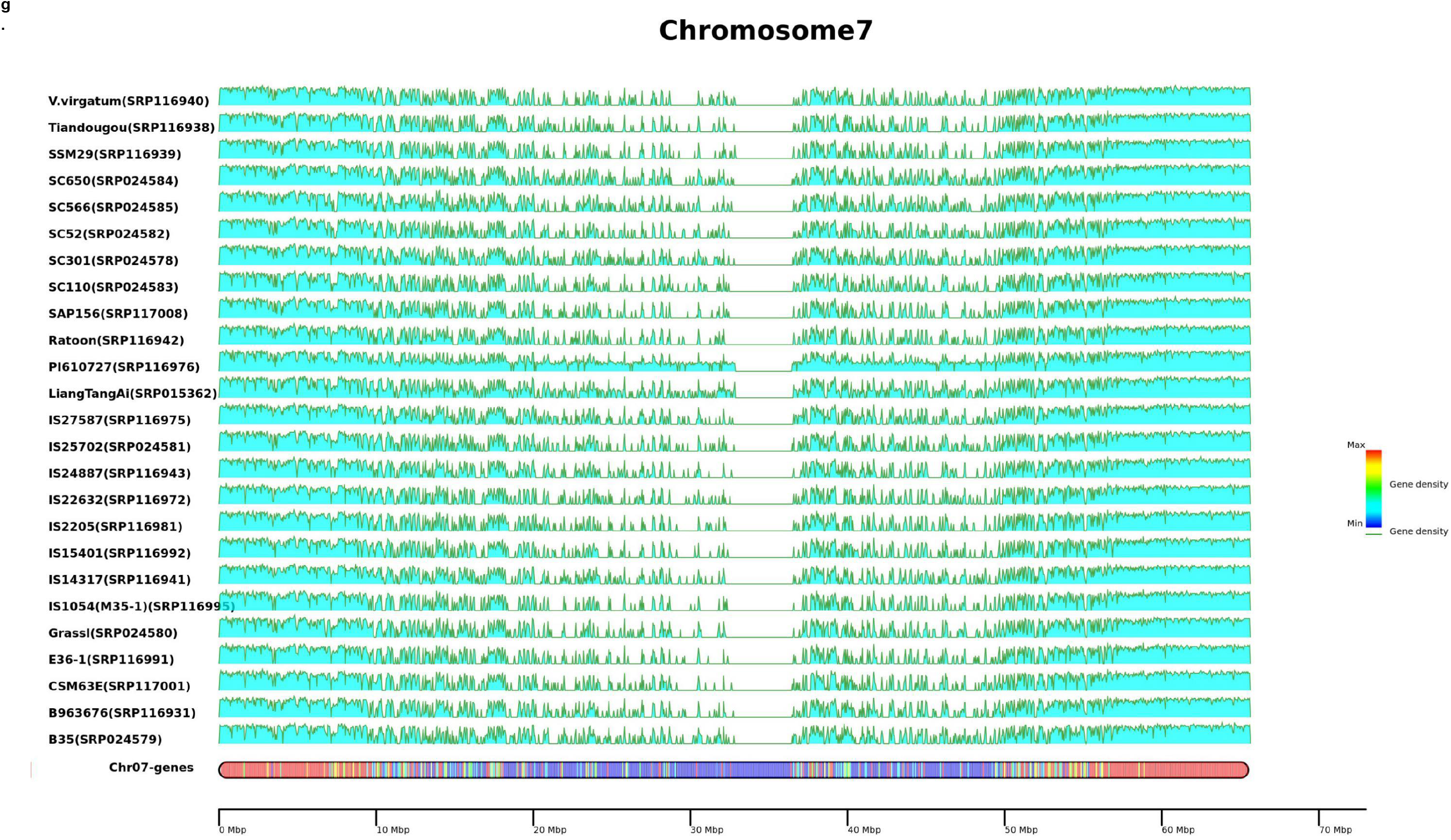
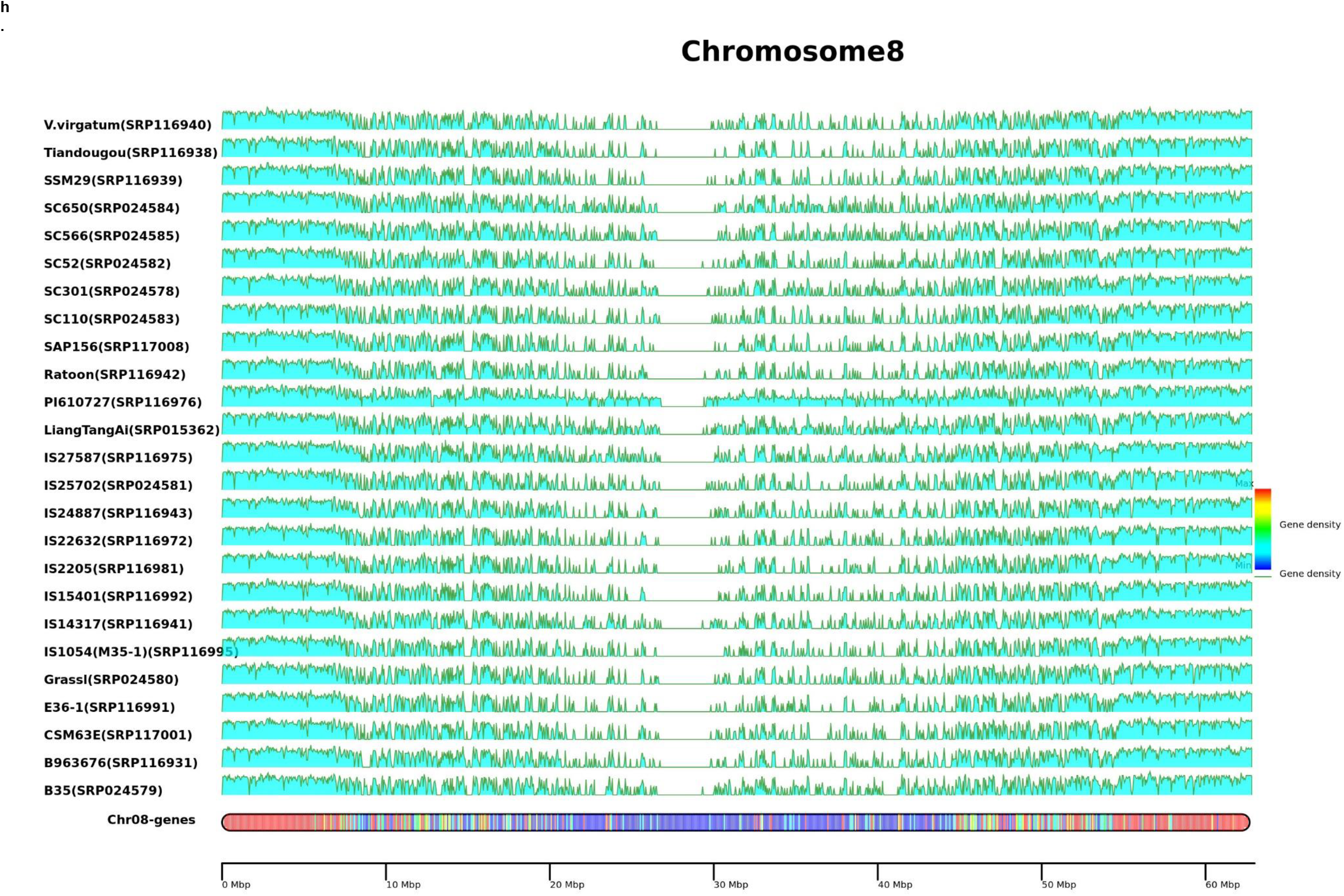
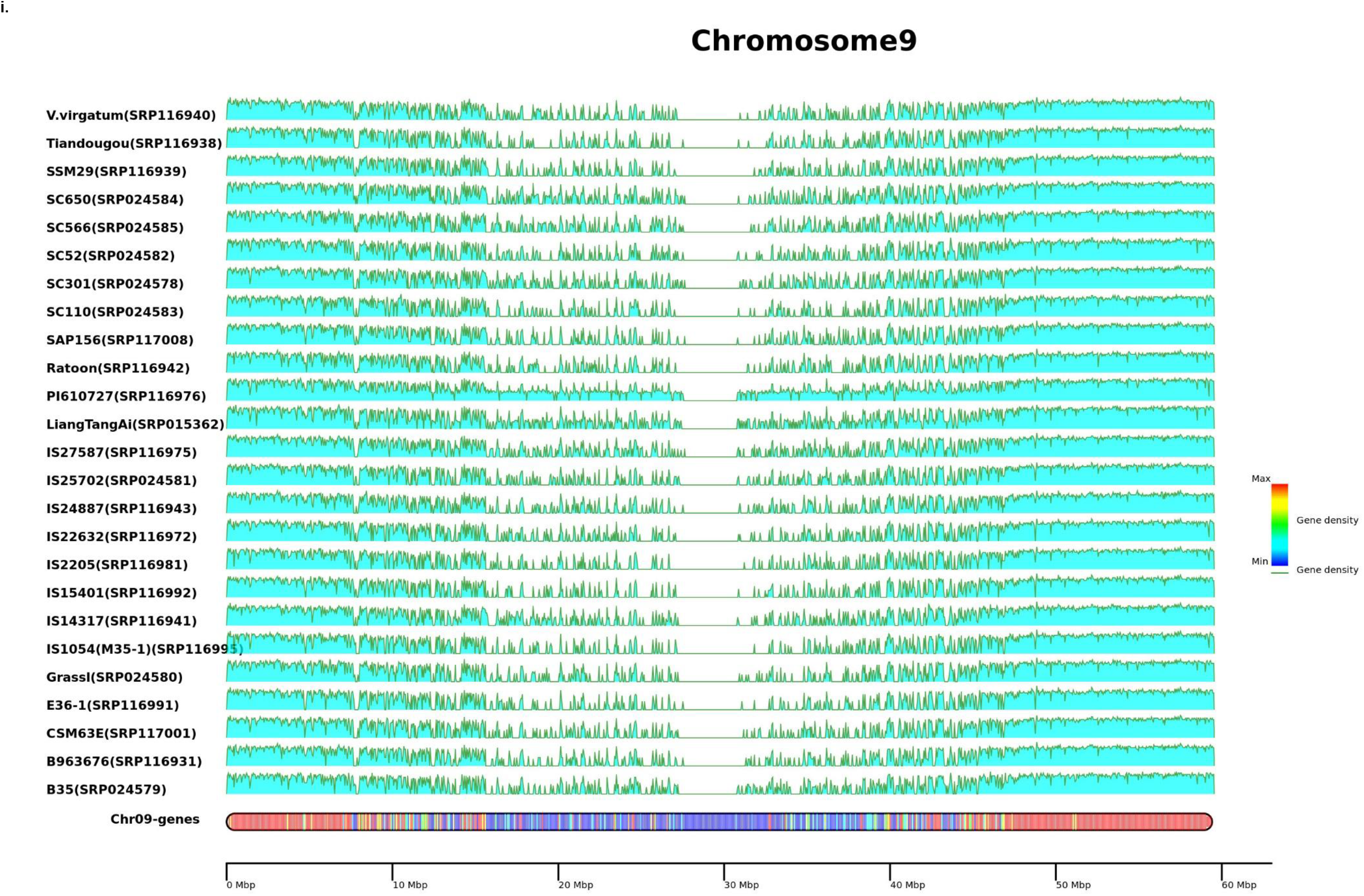
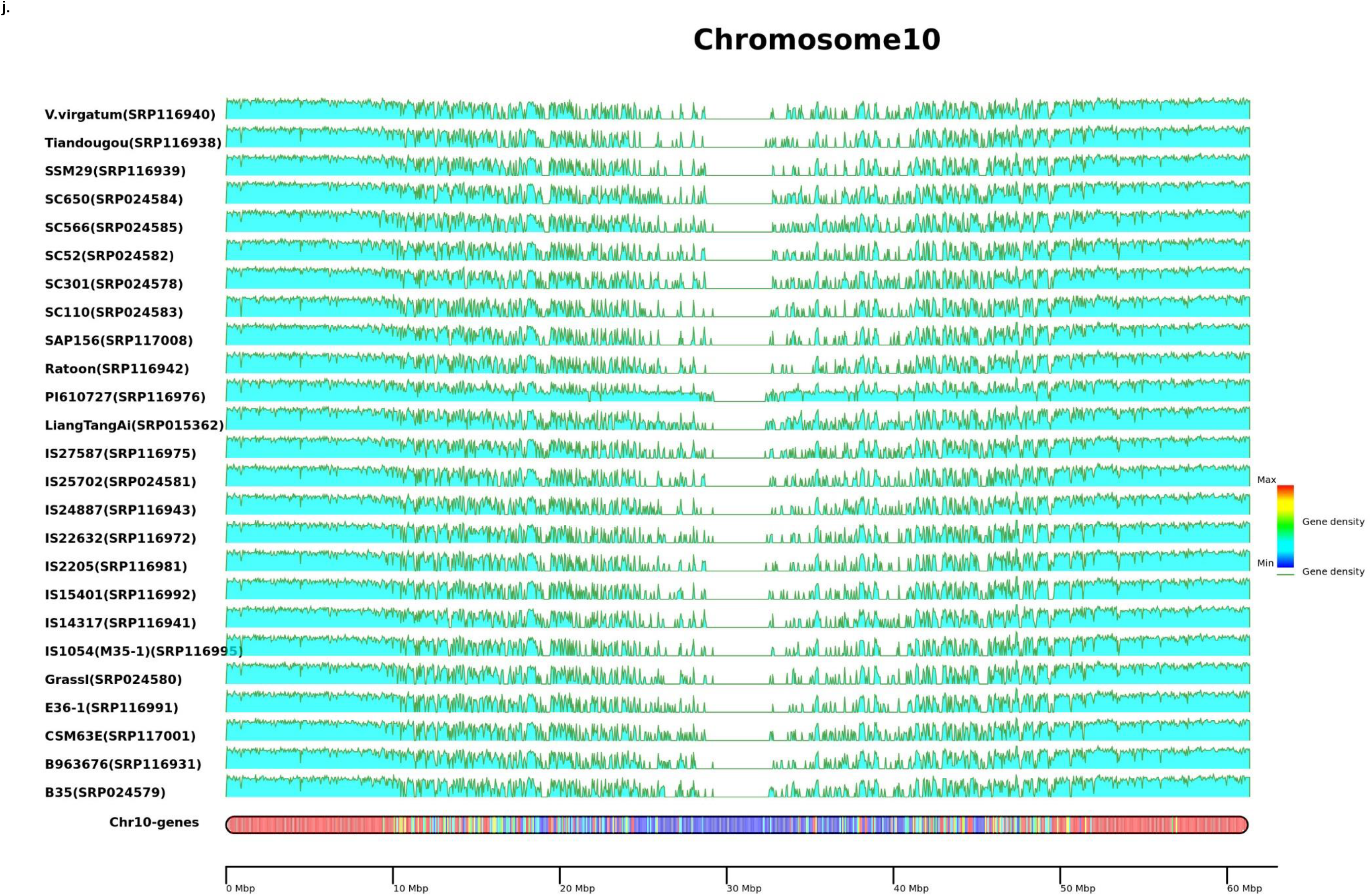
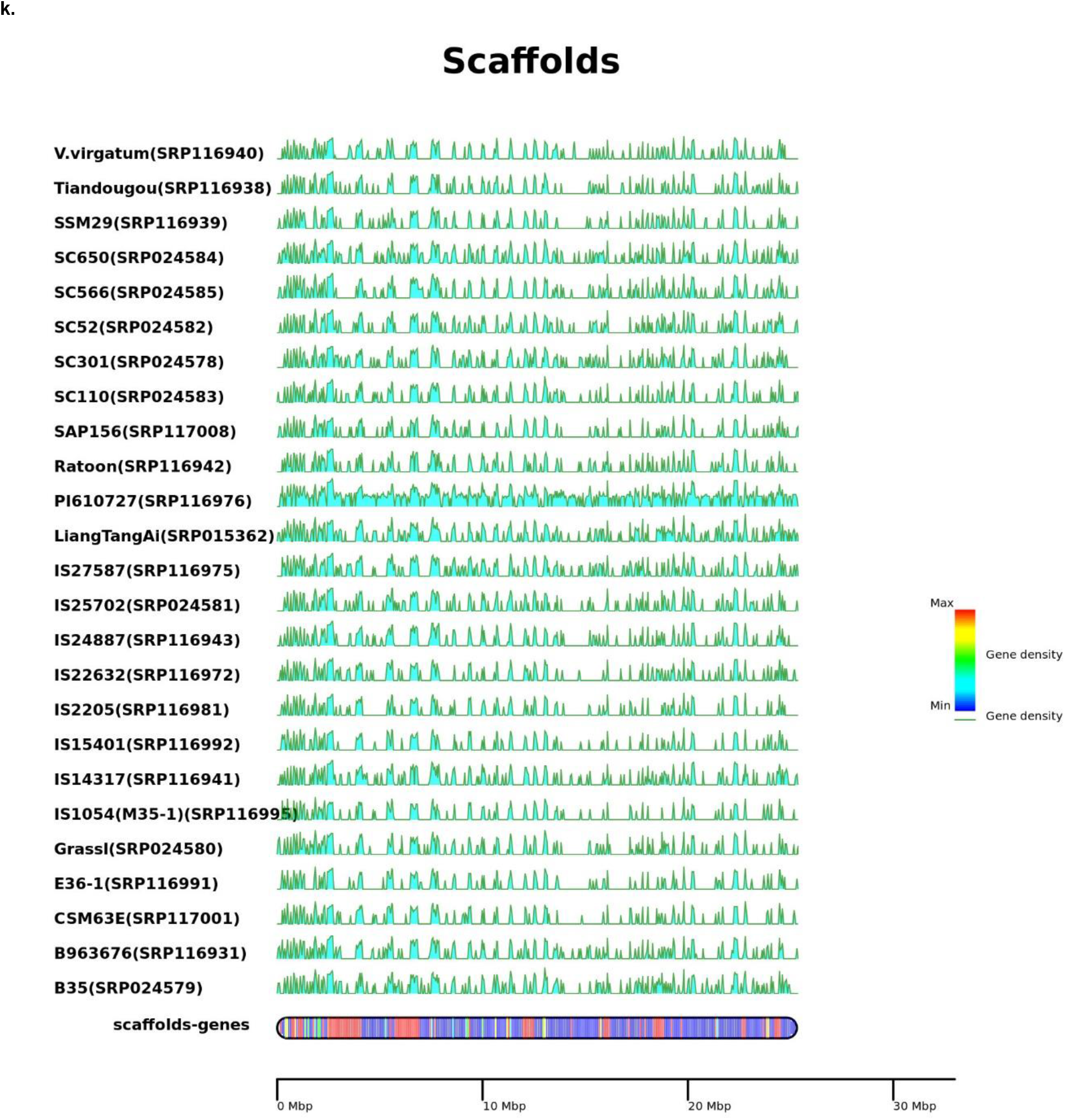
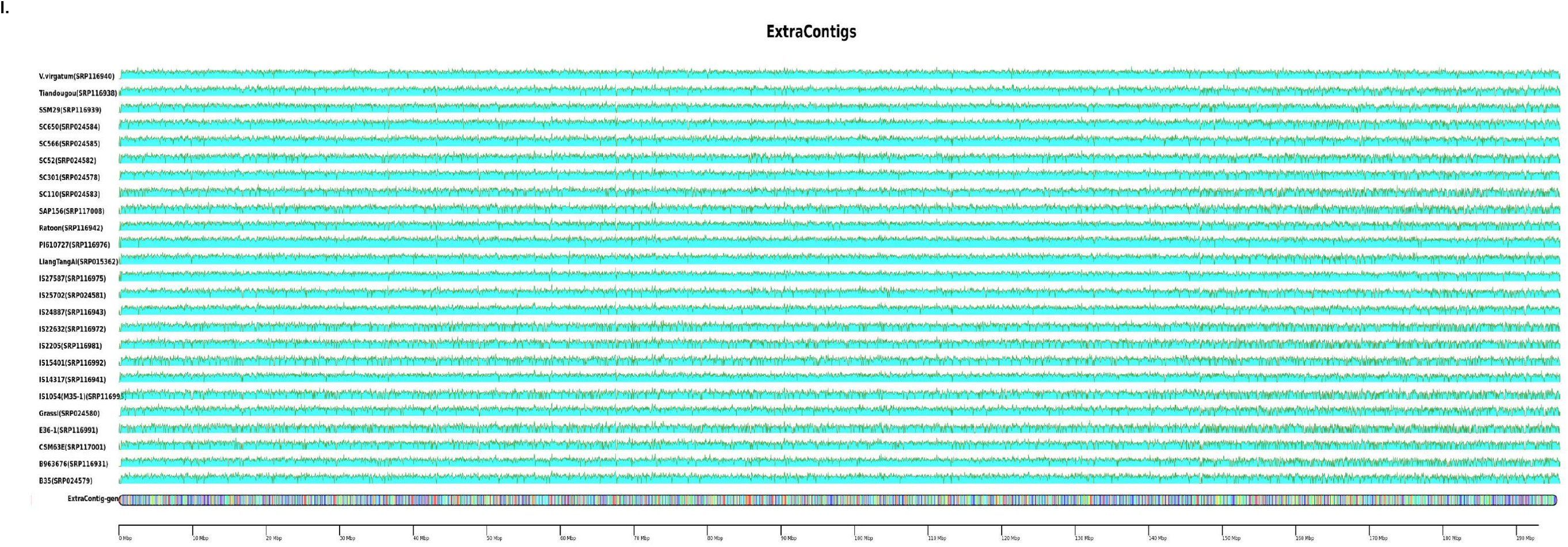
25 accessions (names and NCBI accessions) RNASeq read mapping density on pan-genome assembly for a-j) Chromosome 1-10 k) scaffold sequence put together as single sequence and l) non reference sequence assembly contigs from sorghum accessions concatenated to single sequence as extra contig sequence.

